# Habitat correlates of cave-dwelling: A radiation-scale analysis of skin traits and comparative transcriptomics of the *Sinocyclocheilus* cavefish

**DOI:** 10.1101/2023.07.16.549170

**Authors:** Xiayue Luo, Bing Chen, Tingru Mao, Yewei Liu, Jian Yang, Madhava Meegaskumbura

## Abstract

With 78 species, *Sinocyclocheilus* cavefish constitute the largest cavefish radiation in the world. They exhibit remarkable morphological diversity across three habitat types: surface (Surface morphs, Normal-eyed, variably colored), exclusively-cave-dwelling (Stygobitic morphs, Eyeless, depigmented), and intermediate between cave and surface (Stygophilic morphs, Micro-eyed, partially depigmented). Distinctive traits of *Sinocyclocheilus* include variations in eye and skin conditions associated with their habitat, despite the role of the skin in sensing environmental changes, its habitat correlates are less understood, compared to the well-studied eye conditions. Here, we analyzed the correlation between *Sinocyclocheilus* skin morphology and its habitat, utilizing morphological and transcriptomics-based methods. We generated RNA-sequencing data for nine species and integrated those with existing data from five additional species. These 14 species represent the primary clades and major habitats of these cavefish. Data on skin color and scale morphology were generated and 7374 orthologous genes were identified. Using a comparative transcriptomics approach, we identified 1,348 differentially expressed genes (DEGs) in the three morphotypes. GO and KEGG enrichment analyses suggest that these species have evolved different strategies for energy metabolism, immunity, and oxidative stress in different habitats. We also found 329 positive selection genes (PSGs) in the skin of these species that are mainly involved in immunity, apoptosis, and necrosis, indicating potential adaptations to their habitats. The maximum likelihood phylogenetic tree, based on 1369 single-copy orthologous genes of the species, was largely concordant with the currently established RAD-seq and mt-DNA based phylogenies, but with a few exceptions. Species with higher cave dependence present lighter coloration, fewer dark blotches, and diminished scale morphology and coverage. PCA and cluster analysis suggested that cave-dwelling species, characterized by the absence of black blotches, have similar expression patterns, indicating convergence in cave adaptation. Variations in tyrosine metabolism may explain pigmentation differences among species in diverse habitats. Our study highlights the significance of habitat in shaping skin metabolism, pigmentation variation, and morphology while offering insights into the molecular mechanisms driving these habitat-specific adaptations in *Sinocyclocheilus*. These findings underscore the transcriptional variation in adapting to diverse environments and contribute to future studies on the evolution and ecology of cavefish.

## INTRODUCTION

Cave-adapted organisms, particularly cavefish, make up a significant portion of vertebrates inhabiting caves. They present unique opportunities for evolutionary biologists to study the patterns of adaptation to new environments (Jeffery, 2001). Over 300 cavefish species have been discovered worldwide, evolving rapidly from their surface-dwelling ancestors, making them ideal for comparative analysis of adaptive processes (Borowsky, 2018, Policarpo et al., 2021, Fortune et al., 2020). The best known among these is the well-studied *Astyanax mexicanus* cavefish system, a single species that includes surface and cave-adapted populations representing two distinct morpho-habitat types (Gross, 2012, Loomis et al., 2019, Krishnan et al., 2020). Interestingly, phylogenetically distant groups display convergent stygomorphic traits, providing opportunities for cave-adaptation related comparative studies (Romero and Paulson, 2001, Protas and Jeffery, 2012, Stahl and Gross, 2017).

China’s *Sinocyclocheilus* cavefish, with 78 species, constitute the largest cavefish radiation in the world (Wen et al., 2022, Mao et al., 2022b, Luo et al., 2023, Xu et al., 2023). Phylogenetic analyses based on mitochondrial and nuclear DNA have consistently resolved *Sinocyclocheilus* as a monophyletic genus, comprising of 4-6 main clades, that share a common normal-eyed surface dwelling ancestor (Zhao and Zhang, 2009, Jiang et al., 2019, Mao et al., 2021). *Sinocyclocheilu*s genus can be grouped into three categories based on habitat occupation: Surface (living outside caves, SU), Stygophilic (cave-associated habit, SP), and Stygobitic (exclusively cave-dwelling habit, SB) species (Yang et al., 2016, Zhao et al., 2021, Zhou et al., 2022b). The eye morphology and skin coloration of these fish strongly correlate with their habitats and hence, eye-condition can be used as a proxy to identify their habitat associations (Jiang et al., 2019, Zhao et al., 2021, Mao et al., 2021). SUs have normal eyes and yellow or charcoal gray coloration, while SBs lack eyes and have white-pink, translucent skin (Li et al., 2020, Mao et al., 2021). SPs usually have micro-eyes and variable coloration, although there are exceptions (Mao et al., 2021, Chen et al., 2022). There are several instances of independent evolution of SUs (in two clades) and SBs (Three Clades) within the diversification (Wen et al., 2022, Mao et al., 2022b). So far, *Sinocyclocheilus* research has predominantly focused on eye regression, leaving the skin relatively understudied (Meng et al., 2013, Huang et al., 2019, Zhao et al., 2021).

Epidermal adaptations are essential for organisms conquering new environments (Jablonski and Chaplin, 2010, Ángeles Esteban, 2012). *Sinocyclocheilus* have repeatedly evolved both regressive and constructive skin traits as adaptation to cave environments. Regressive traits include: reduced pigmentation, reduction in scales and constructive features include increased fat accumulation and enhanced non-visual sensory abilities such as well-developed lateral lines and neuromast systems (Yang et al., 2016, Yoshizawa et al., 2014, Chen et al. 2022). Hence, the independent evolution of the morphotypes of *Sinocyclocheilu*s provide an attractive system to investigate the genetic basis of convergent adaptations of their skin at the molecular level.

Transcriptomics based tools provide an unprecedented opportunity to understand the gene expression profiles and patterns of genetic variation behind the independent evolution of these diverse cavefish morphotypes. At the broader scale, studies on *Astyanax mexicanus* cavefish and *Sinocyclocheilus* cavefish have shown how protein sequence alterations (transcriptome) play a role in eye morphology and color degeneration in cavefish (Hinaux et al., 2013, McGaugh et al., 2014, Torres-Paz et al., 2018, Huang et al., 2019, Zhao et al., 2020, Li et al., 2020). However, little is known about the mechanisms of scale degradation (Simon et al., 2017, Yang et al., 2016). The convergent evolution of skin coloration is largely driven by the similar sensory adaptations to similar light environments that evolved independently in each species (Moran et al., 2023). However, the relationship between scales and cave habitat is unclear (Zhao and Zhang, 2009). Therefore, it is also necessary to explore the morphological traits of the skin in an evolutionary comparative framework.

It is known that the habitat influences skin immunity, microbial composition, and host skin sensitivity (Scharsack et al., 2007, Zhou et al., 2022b, Peuß et al., 2020). The maintenance of the health of an organism is a result of a dynamic interplay between the microbiota, host skin cells, and immune system, which work synergistically in a mutually beneficial manner (Belkaid and Hand, 2014, Austin, 2006, Ellison et al., 2021). In a study involving the three representative morpho-species, it has been shown that *S. rhinoceros* (SP) displayed the strongest innate immunity, which suggests this as a possible adaptation for greater habitat heterogeneity (Yang et al., 2016).

Caves present a challenging environment characterized by limited food resources and low diversity (Gibert and Deharveng, 2002, Lafferty and K., 2012). Under such conditions, skin functions may be significantly impacted, with effects observed in mucus composition and immune response. As a result, there may be an increased risk of harmful microbial infections and associated health-related issues (McCormick and Larson, 2008).

Multispecies transcriptomics offer new insights into the origins of adaptive phenotypes in cavefish (Stahl and Gross, 2017, Meng et al., 2018, Qi et al., 2018). It can also reveal plasticity or adaptive changes in habitats of related genes that may mediate energy metabolism (Riddle et al., 2018, Lam et al., 2022), immune regulation (Peuß et al., 2020, Huang et al., 2016), and oxidative stress (Krishnan et al., 2020). However, the broader extent of such responses in *Sinocyclocheilus* skin remains underexplored, especially regarding the diverse morphological variations and intriguing immune mechanisms observed in SPs.

To better understand the habitat correlations, morphotypes, and genetic basis of *Sinocyclocheilus* cavefish skin related adaptation, we conducted a comprehensive radiation-scale analysis. These incorporated representatives from the major clades and the three main habitat types. We envision that this approach will allow us to uncover subtle patterns pertaining to the complex interplay among gene regulation, physiological adaptations, and the unique constraints imposed by different habitats on the skin of these cavefish. Specifically, we focus on the following objectives: (1) investigating the shared and derived adaptive mechanisms of the skin of cavefish to diverse habitats; (2) examining the phylogenetic relationships among species based on transcriptomic analysis of their skin-related genes; and (3) elucidating the associations between variations in skin color, scale morphology, and their respective habitats, by integrating molecular mechanisms with observations of morphological traits of the skin.

## MATERIALS & METHODS

### Sample collection

Animal care and experimental protocols were approved for this study by the Guangxi University Ethical Committee under the approval document (GXU-2021-126). To investigate the adaptive mechanism for cave-dwelling in the *Sinocyclocheilus* radiation, 9 species were collected from caves in Yunnan, Guizhou and Guangxi of China during 2019 – 2021 as a part of an ongoing phylogenomic study. Cavefish were transported alive to the laboratory in oxygenated plastic bags in a cooler box to both keep the temperatures low (18 – 20℃) and to provide darkness. Following morphological observations (mentioned below), they were anesthetized using 30 mg/L MS-222 (ethyl 3-aminobenzoate methanesulfonate; Sigma-Aldrich). The skin tissue was taken from the right side near the back of each individual in approximately 0.8 × 1.0 cm under sterile conditions. Three biological repeats were taken for each species. The tissue extraction was done in a DNA/RNA-free clean room. Biopsied tissues were placed in RNAlater and stored in an ultra-low temperature freezer at -83℃ prior to further analysis.

Given their rarity, sampling difficulty, and the need for a representation of the diversification, we enhanced taxon sampling by adding data for five additional *Sinocyclocheilus* species from previous studies (details provided below). Overall, the dataset included species representing the four main clades in the context of Mao et al. (Mao et al., 2021), as well as the three main eye-types / habitats (Normal-eyed SU, Normal-eyed SP, Micro-eyed SP, and Eyeless SB), representative of the *Sinocyclocheilus* radiation (Figure 1, Supplementary Table S1). The eye-type in this study serves as a practical, easily observable feature that corresponds with the habitat occupation of these species.

**Figure 1.**
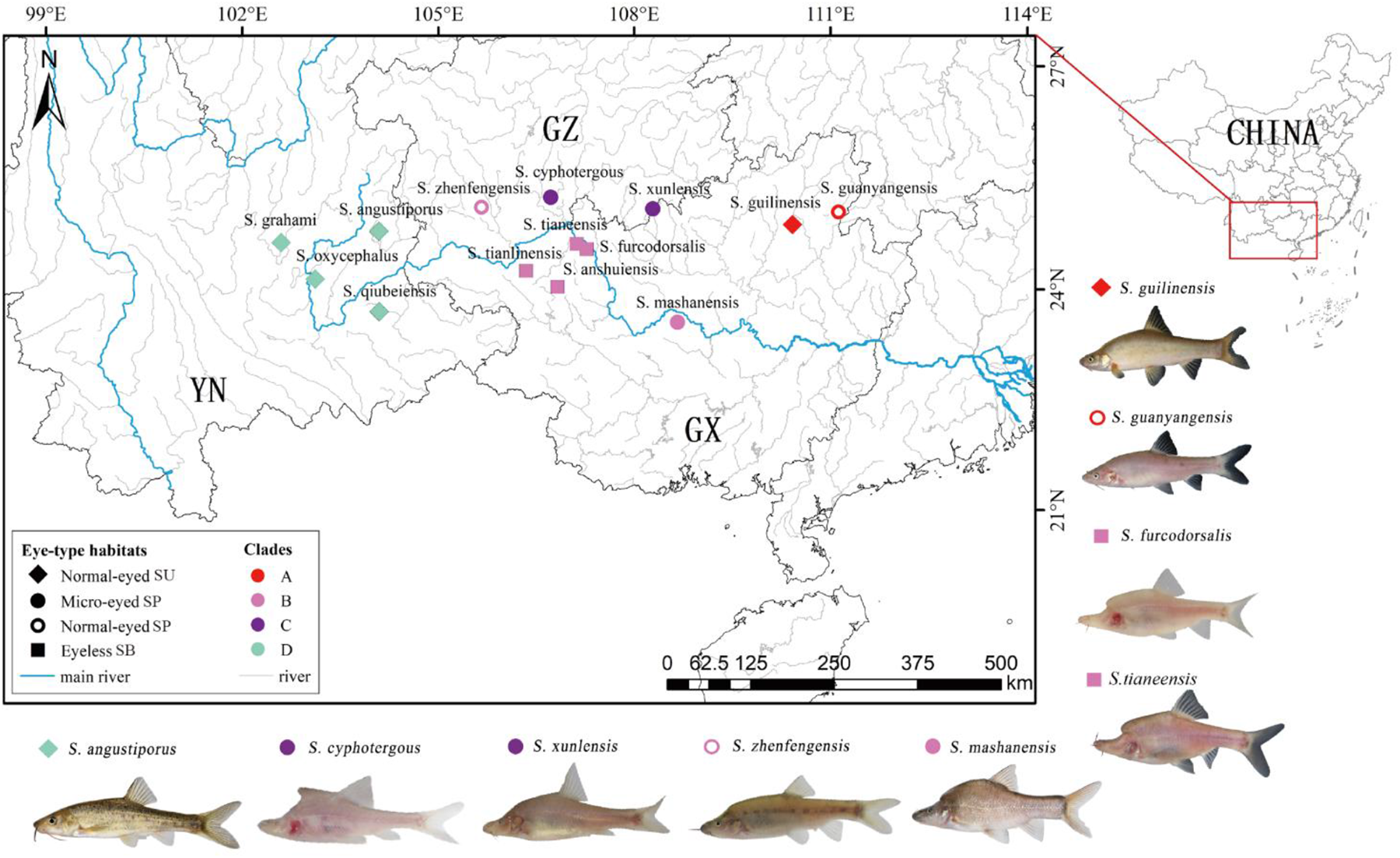
Geographic distribution of sampling sites, morphotypes and clade affiliations for the 14 species representative of the 4 major clades of the *Sinocyclocheilus* radiation. Symbols represent eye-types and habitats; color represents their clades.

### RNA extraction and transcriptome sequencing

In order to generate transcription data for *Sinocyclocheilus* skin from the morphotypes, the total RNA was extracted from the 27 skin samples (9 species that we sampled) using Trizol methods; mRNA was captured from total RNA through Oligo (dT). To synthesize cDNA, fragmented mRNA was utilized as a template in the M-MuLV reverse transcriptase system. Random oligonucleotides acted as primers to initiate the synthesis of the first cDNA strand. RNaseH was then used to degrade the RNA strand, and dNTPs were added under the DNA polymerase I system to synthesize the second cDNA strand. The library was constructed using the NEBNext® Ultra™ RNA Library Prep Kit (Illumina, USA). The RNA quality and concentration were measured using an Agilent Bioanalyzer 2100 (Agilent Technologies, Santa Clara, CA, USA) and Qubit® RNA Assay Kit in Qubit® 2.0 Fluorometer (Life Technologies, Carlsbad, CA, USA). The effective concentration of the library was again accurately quantified by qPCR to ensure the quality of the library. An Illumina NovaSeq 6000 was used to sequence the library, producing paired-end reads of 150 bp each. The library construction and sequencing were conducted at Beijing Novogene Bioinformatics Technology Co., Ltd.

### Data preprocessing

In addition to our data, RNA-seq data for 5 species were obtained from two public databases. We downloaded transcriptome datasets for 3 species from the public DRYAD database: *S. oxycephalus*, *S. tianlinensis*, and *S. qiubeiensis* (https://datadryad.org/stash/share/t5cZIXoVUgyhpzEP6z-GN6xjc5EU3TvPPEwbdIo7siI), and transcriptome datasets for 2 species: *S. grahami* (NCBI: SRR2960332) and *S. anshuiensis* (NCBI: SRR2960751) from the NCBI database. The skin transcriptomes from the 9 species from our study and the 5 additional species were evaluated with FastQC v0.11.9 (Andrews, 2010). To ensure the accuracy and reliability of our data, we implemented a rigorous filtering process. This involved removing reads containing adapters, poly-N sequences, and those with low-quality scores. The resulting high-quality, clean reads formed the basis for all downstream analyses.

### Transcriptome assembly, annotation and selection of orthogroups

We used Trinity v2.8.6 (Grabherr et al., 2011) to assemble clean reads for each species and extracted the longest transcripts (unigenes), to obtain single gene sequences. Subsequently, the CD-HIT v4.6.8 (Fu et al., 2012) was used with a 95% threshold to cluster sequences and eliminate redundancy in the final assembly. To predict the full open reading frames (ORFs) for each gene, we used TransDecoder (http://transdecoder.github.io/) default parameters. We then annotated the resulting protein sequences using BLASTP (Blast+ v2.6.0) using the UniProt database (v2022_05) (Mahram and Herbordt, 2015). We retained only UniProt germplasm from the top 10 hits for each species and selected the final UniProt ID based on the number of species that used it as the best match. Using OrthoFinder v1.1.2. (Emms and Kelly, 2015), we performed gene clustering while retaining only those containing at least one transcript per species. These filtering steps resulted in 30,923 annotated orthologous groups for downstream analysis of expression differences.

### Construction of phylogenetic trees and screening of positive selection gene (PSG)

For phylogenetic inference, we utilized single-copy gene families obtained from OrthoFinder, as previously described. We discarded gene families comprising sequences shorter than 200 amino acids. The retained amino acid sequence families were aligned with Muscle 5 (Edgar, 2022), with default parameters. To excise ambiguously aligned regions, we used Gblock v0.91b (Castresana, 2000). We then conducted a maximum likelihood (ML) phylogenetic analysis on the refined set of 1,369 single-copy orthogroups using IQ-TREE 2 to construct an unrooted phylogeny; a bootstrap analysis was performed to determine node support (Minh et al., 2020). The resulting ML phylogenetic tree was visualized using Figtree (Rambaut, 2009).

To understand the functional genes that may facilitate the adaptation of *Sinocyclocheilus* to its environment, orthologous of the skin of 14 species were tested for signals of positive selection. The dN, dS, and dN/dS values of the orthologous were calculated using the CodeML program of the PAML package (Yang, 2007). The CodeML parameter was set to “Runmode = 0, Model = 0”. Then, ω> 1 can be judged to have experienced positive selection pressure effects in this gene, and a total of 329 genes were screened. KEGG enrichment of PSGs was performed using KOBAS v2.0.12, and the PSGs in significantly enriched pathways were annotated based on the Evolutionary Genealogy of Genes: Nonsupervised Orthologous Groups (eggNog) database (Xie et al., 2011, Huerta-Cepas et al., 2016).

### Differentially expressed gene (DEG) analysis of orthogroups

To study how orthologous genes are expressed in three different habitats – SB, SP, and SU – we conducted a mapping analysis of high-quality reads of each species against its corresponding representative orthogroups obtained earlier; for this, we used Bowtie 2 v2.2.9 for direct homology filtering (default settings) (Langmead and Salzberg, 2012), and for gene expression levels analysis we used RSEM v1.2.26 (default setting) (Li and Dewey, 2011). To analyze the gene expression profiles of each species, we utilized the ggbiplot package to conduct a principal component analysis (PCA) following the method outlined by Yeung and Ruzzo (2001). The ggplot2 package was used to statistically generate stacked bar graphs with clustered trees reflecting the similarity between species and habitat groups, as well as information on the expression profiles of the orthologous genes for each species (Thiergart et al., 2020). We then identified differential expression in the orthogroups between species using edgeR and adjusted the resulting P-values through the application of the Benjamini and Hochberg method, which effectively estimates the false discovery rate (FDR). We considered differentially expressed genes to be those with absolute log2 fold change (|log2FC|) greater than 1 and FDR less than 0.05. These thresholds were chosen to determine statistical significance. Finally, GOseq and KOBAS v2.0.12 were used to perform GO enrichment analysis and KEGG enrichment analysis of differentially expressed orthogroups (Kanehisa et al., 2008, Young et al., 2010). P-values were obtained using a hypergeometric test and the significance term with an adjusted P-value threshold of 0.05. Using these methods, we identified significantly enriched biological pathways, including important biochemical metabolic and signal transduction pathways across the orthogroups.

### Skin photography

Using a digital camera (Canon EOS 6D Mark II AF-A) set at a fixed distance of 0.3m from the tank and LUX = 45 for ambient light, we captured images with the following settings: Shutter speed: 1/250s; F/20; ISO 200. Next, the fish were returned to their aquarium system (pH: 7.0-8.0; temperature: 19 ± 1℃; dissolved oxygen: 8.5 mg/L) devoid of light to maintain their natural skin characteristics. Subsequently, we examined the pigmentation and scales in samples preserved in 70% ethanol using a Leica M165FC stereomicroscope.

### Calculation of mean color, dark clustering and measurements of scales

To compare the body color differences of the *Sinocyclocheilus* genus, we used the Color Summarizer v0.8 (http://mkweb.bcgsc.ca/colorsummarizer/analyze) to digitize the average color values of the skin. In order to ascertain the average red-green-blue (RGB) color component of the image, we employed a systematic approach. Firstly, a set of reference colors that had the least deviation from the image colors were identified, which allowed to discern variation in color with greater precision. Specifically, 5 reference colors denoted by “k” were selected. Subsequently, 5 representative color ratios from each sample taken. Finally, the average RGB color values were calculated from these ratios to obtain an overall RGB color component of the image. This procedure provided an accurate and reliable estimation of the RGB color values. This process was repeated three times separately for the three samples collected to obtain a representative color for each *Sinocyclocheilus* species. Our analysis focused on the 5 most representative color ratios in order to calculate the proportion of darker blotches. For instance, in a particular species, mean color values of RGB (250, 250, 250) were observed in the white areas while the darker black areas had color values closer to RGB (0, 0, 0). We postulated that a higher concentration of melanin deposition is associated with a decrease in RGB values towards RGB (0, 0, 0). The area and number of lateral line scales were counted for each sample using ImageJ (Abràmoff et al., 2004). The scale sizes were classified into three categories: no-scales, small-scales, and large-scales, and the degree of scale cover was classified as full-cover, no-cover, and partial-cover.

## RESULTS

### Transcriptome sequencing data and identification of orthologous

To understand the molecular mechanisms underlying skin coloration in *Sinocyclocheilus* species, we generated high-quality transcriptome sequencing data from skin tissue samples of 9 distinct *Sinocyclocheilus* species (Supplementary Table S2). Following this, a *de novo* assembly was performed for sequences from the 14 species. Among them, *S. oxycephalus* (205,993) has the highest number of transcriptomes and *S. grahami* (80,118) has the lowest. The average transcript length post-assembly varied between 856 and 1,158 bps, while the N50 length ranged from 1,621 to 2,414 bps. Upon removal of redundancy, a total of 384,146 genes were identified within orthogroups. Overall, orthogroups accounted for 94.3% of the genes. To facilitate comparative analysis among the various species, we focused on the 7,374 orthologous genes present in all examined species. These orthologous gene families were clustered, and 1369 single-copy orthologues genes were obtained after multiple sequence alignment and low quality pairwise pruning. These orthologous genes were subjected to quantitative comparative analysis.

### Distinct habitats influence gene expression variation

A total of 1,348 differentially expressed genes (DEGs) were identified on the skins of *Sinocyclocheilus* species by conducting a pairwise comparison across the three habitats (Figure 2, Supplementary Table S3). When compared to SUs, SPs exhibited 200 up-regulated and 467 down-regulated genes; SBs displayed 292 up-regulated and 394 down-regulated genes. The majority of DEGs were up-regulated in SUs and the highest number of down-regulated DEGs was observed in SPs (Figure 2A). We also found more unique gene up-regulation in SP vs. SB, and that more of their shared genes were consistently down-regulated compared to SUs (SU vs. SP and SU vs. SB, Figure 2B). This suggested a significant impact of cave habitats on gene expression patterns.

**Figure 2.**
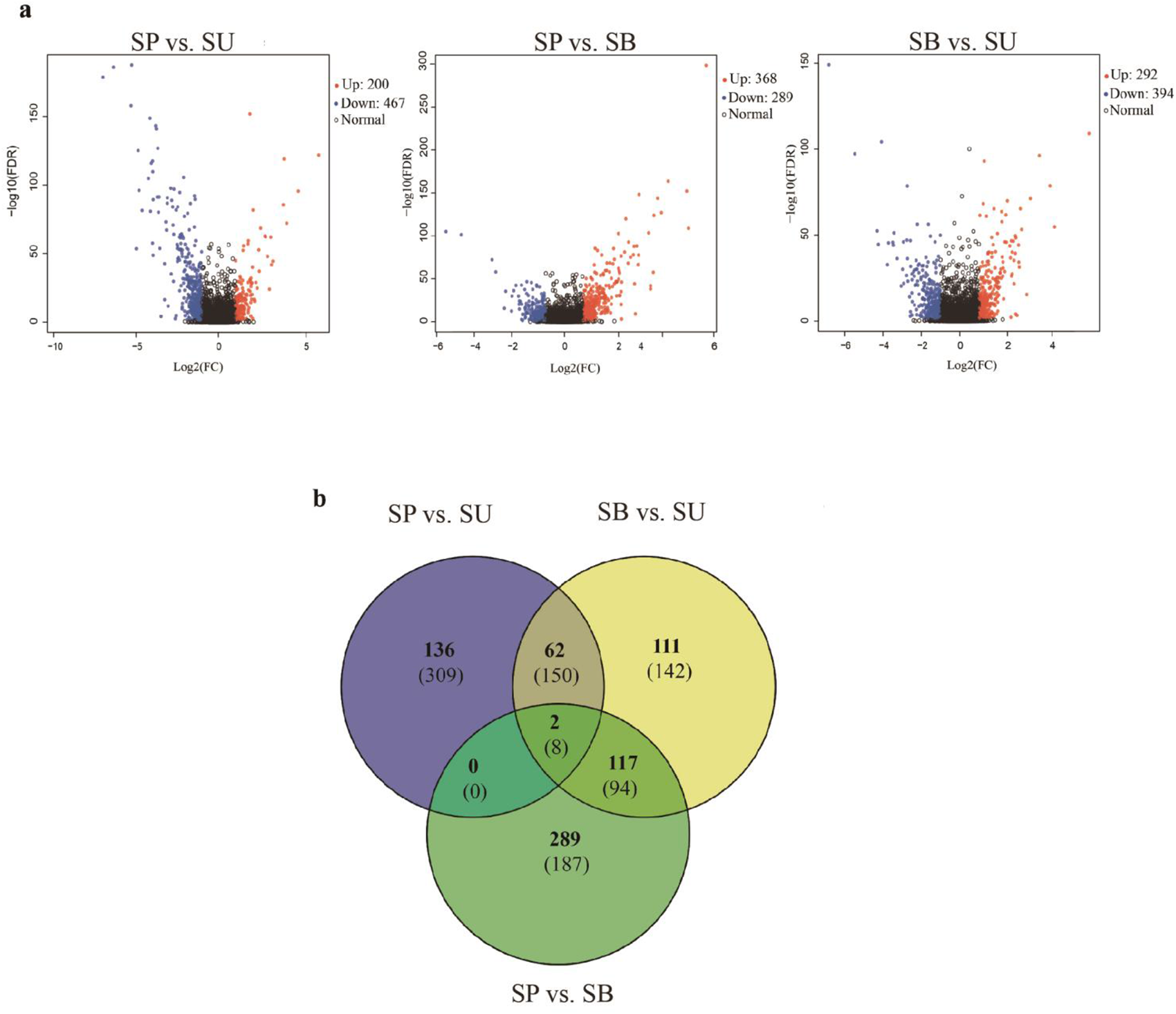
(A) Volcano plots of the distribution of DEGs between SU vs. SP, SP vs. SB and SU vs. SB. The x-axis shows log 2-fold change in gene expression. The y-axis shows -log 10 (p-value). The further away from 0 on the x-axis, the greater the change in expression, and the higher the y-axis, the greater the significance. Blue dots indicate up-regulation, red dots indicate down-regulation and black dots indicate no change in expression in the DEG. (B) Venn diagrams depict shared and unique variations in gene expression among the three main habitats (SU, SP and SB). The numbers in each section correspond to the number of DEGs from gene expression estimates. The number of up-regulated DEGs is listed at the top (in bold) and the number of down-regulated DEGs is listed at the bottom.

GO enrichment analysis of 1,348 DEGs in different eye-types/ habitats identified 630, 565 and 799 significantly enriched GO terms in SU vs. SP, SP vs. SB and SU vs. SB, respectively (*P* _ajd_ <0.05) (Supplementary Table S4). These DEGs were involved in functional responses mainly related to stimulus responses, catalytic activity, multi-organism process, and immune system process (Figure 3A). We found that the co-enriched terms in these three groups were energy metabolism related (Supplementary Table S5). We also found two major categories of GO terms have been enriched in SU vs. cave dwellers (SP and SB). Stimulus response-related terms such as external biotic stimulus, bacterial, fungal, and viral reactions, and numerous immune-related terms. SU vs. SP was mainly related to cellular immunity, such as leukocyte mediated immunity (GO:0002444), regulation of macrophage derived foam cell differentiation (GO:0010743); while SU vs. SB increased the regulation of apoptosis, such as: positive regulation of MAPK cascade (GO:0043410), regulation of neuron apoptotic process (GO:0043523), regulation of epithelial cell apoptotic process (GO:1904035). Anyhow, this suggested that these observations may be due to the differences in the exogenous biological stimulation of surface water environment. Cave species seemed to have different immune strategies to these external stimuli.

**Figure 3.**
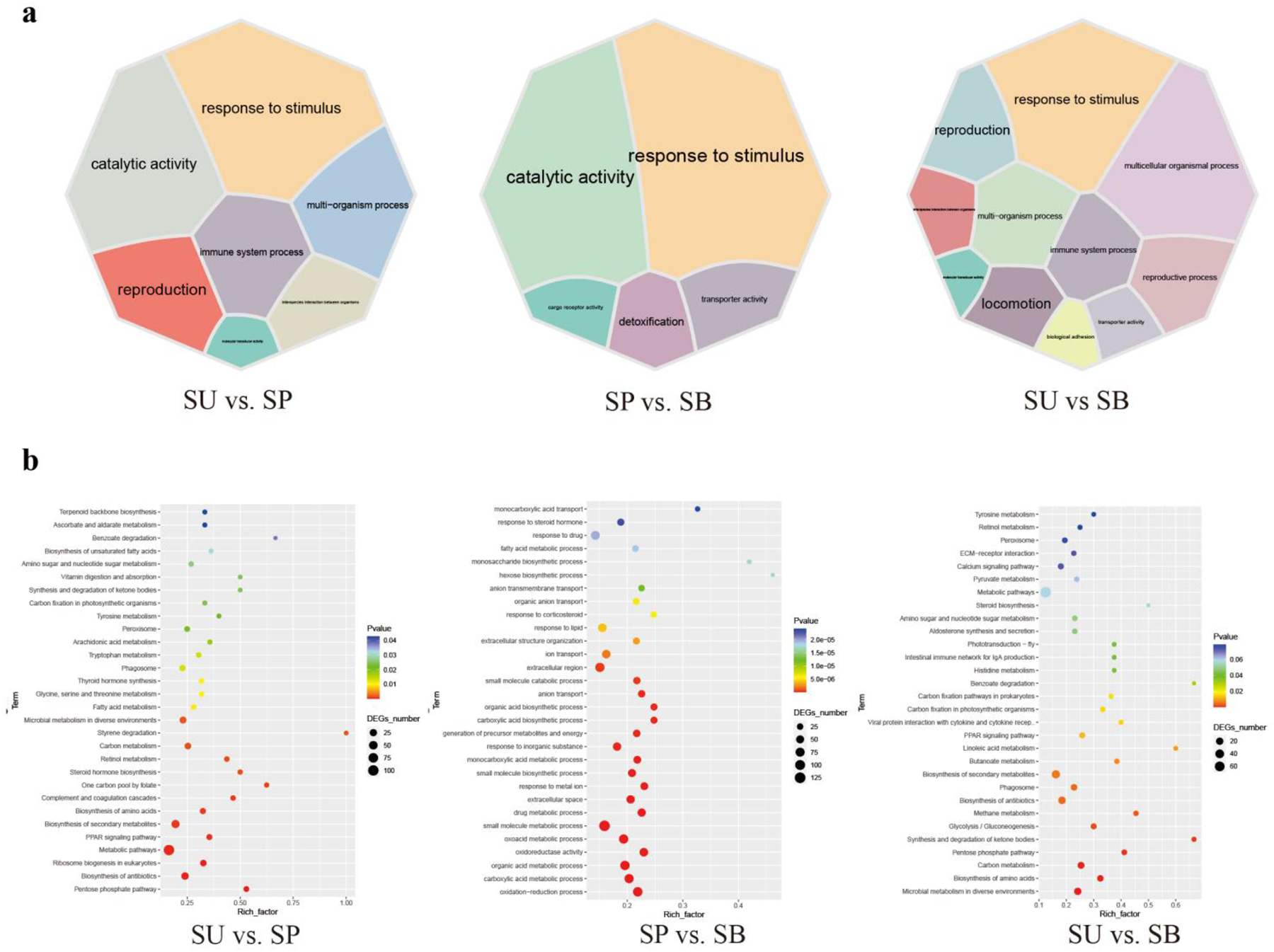
Enrichment maps for DEG among the three main habitats (SU, SP and SB). (A) Classification of GO trems significantly enriched in differentially expressed genes (DEGs). 7 GO categories in SU vs. SP; 5 GO categories in SP vs. SB, and 11 GO categories in SU vs. SB. Different GO categories are displayed in different colors, and the size of the module represents the number of DEGs associated with the corresponding functional item. (B) The top 30 KEGG pathways that are enriched in differentially expressed genes (DEGs), including SU vs. SP, SP vs. SB and SU vs. SB.

Genes associated with changes in oxygen levels showed differences in expression across different habitats, for example, response to hypoxia (BP), response to decreased oxygen levels (GO:0036293), cellular response to hypoxia (GO:0071456), etc. Interestingly, only in SPs and SBs can respiratory electron transport chain (GO:0022904), mitochondrial respirasome (GO:0005746), respiratory electron transport chain (GO:0022904), and mitochondrial respirasome (GO:0005746) be identified. respiratory chain complex (GO:0098803) and response to pH (GO:0009268) (Supplementary Figure S1A, supplementary Table S5). This may reflect the different cellular and mitochondrial respiration efficiency of cave dwellers.

The number of significantly enriched pathways in three groups was 30, 38, and 22, respectively, and the same pathways were mainly related to metabolisms, such as Pentose phosphate pathway (ko00030), Biosynthesis of amino acids (ko01130), Carbon metabolism (ko01200), PPAR signaling pathway (ko03320) and Microbial metabolism in diverse environments (ko01120), etc. Secondly, immune response-related pathways were Phagosome (ko04145) (*P* _ajd_ <0.05) (Figure 3B, Supplementary Table S6). This suggested that the strong influence of the habitat environment leads to differences in skin metabolism and skin microbial metabolism among different habitat populations. Compared to SPs, all DEGs (28) enriched in the PPAR signaling pathway and Carbon metabolism pathway were up-regulated in SUs, followed by 16 DGEs in SBs. However, all DEGs (9) enriched Oxidative phosphorylation (OXPHOS, ko00190) pathway and 5 DEGs enriched Fatty acid degradation pathway (ko00071) were upregulated in SPs. This may indicate differences in the regulation of energy and mitochondrial metabolism between species in different habitats (Supplementary Table S7). These pathways in different eye-types/habitats showed that there were common molecular mechanisms to understand their habitual differences and evidence of genetic variation and transcriptional plasticity or adaptation in response to environmental change.

In particular, microbial immune-related pathways such as Complement and coagulation cascades (ko04610), Intestinal immune network for IgA production (ko04672), and Viral protein interaction with cytokine and cytokine receptors (ko04061) found in SU vs. cave dwellers (SP and SB) (Supplementary Table S6). This may indicate differences in immune responses to microorganisms between SUs and cave dwellers. Moreover, Hematopoietic cell lineage (ko04640) and ECM-receptor interaction (ko04512), and pathways associated with oxidative stress (OXPHOS, Glutathione metabolism, Cysteine, and methionine metabolism, Metabolism of xenobiotics by cytochrome P450, Drug metabolism – cytochrome P450) exist only in SP vs. SB. This may indicate a widespread oxidative stress response of cave dwellers to stimuli of the cave environments.

We also found that Phenylalanine, tyrosine and tryptophan biosynthesis pathway (ko00400), and especially the Tyrosine metabolism pathway (ko00350) may be influenced by their habitats, and affected melanin differences (Supplementary Table S6).

### Positive Selection Genes in the skin of 14 species

After analyzing the KEGG pathways of these 329 positive selection genes (PSGs), we found that genes under positive selection were most significantly enriched in six metabolic pathways (Table 1). They were associated with viral infection, signaling, and cell necrosis and apoptosis. A total of 18 PSGs were identified in these enriched pathways, among which *the tumor protein p63 regulatory 1* (OG0018161) was under the strongest selection pressure (Supplementary Table S8). These genes may be involved in the adaptation process of *Sinocyclocheilus* to cave dwelling. We also found that *Reticulocalbin 3*, *EF-hand calcium binding domain* (OG0016528), *S100 calcium binding protein U* (OG0016522), and *positive regulation of vitamin D* (OG0016683) have all been implicated in calcium regulation.

**Table 1.**
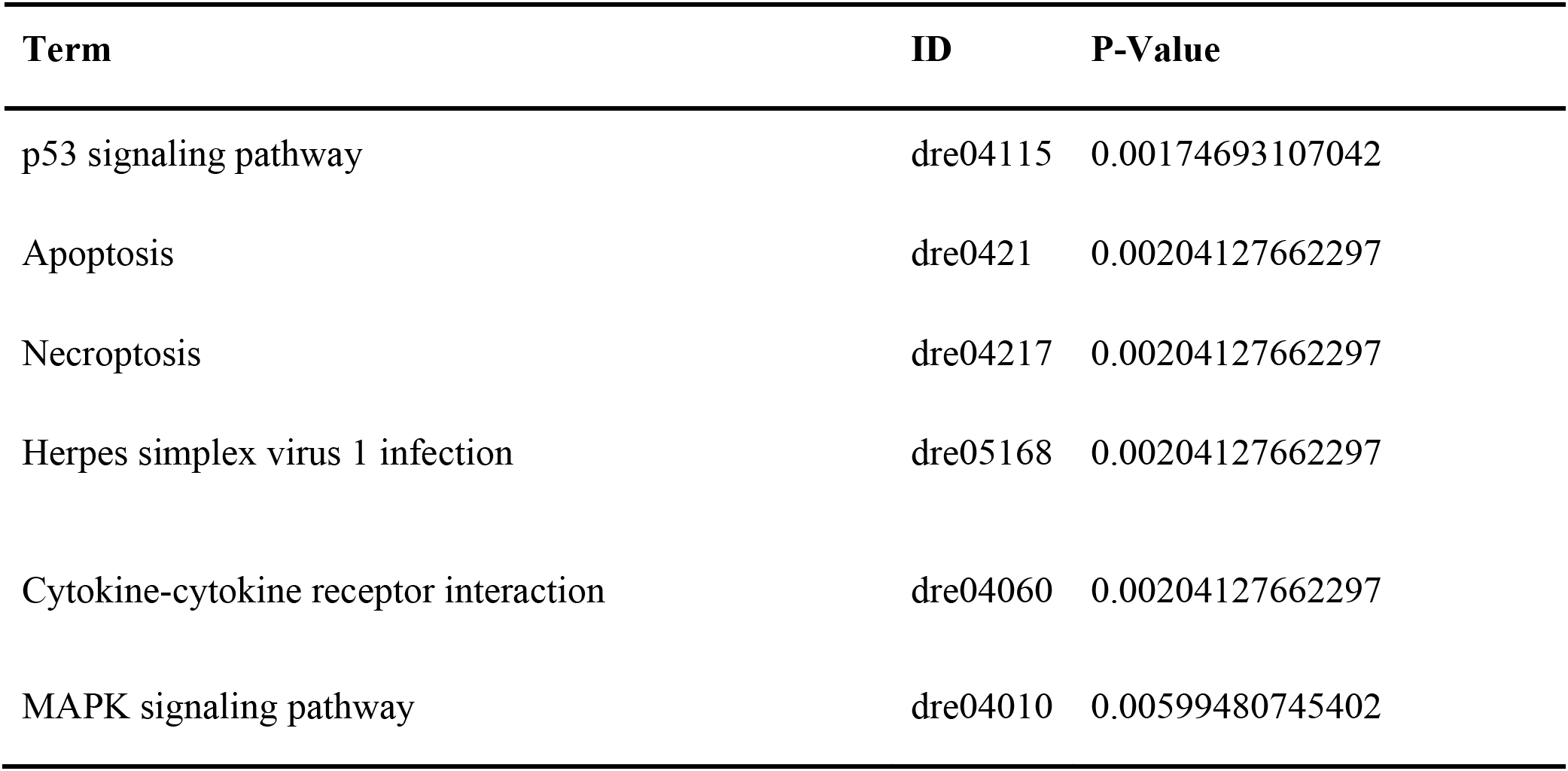
List of positive selection KEGG pathways including their terms, IDs and p-values.

| Term | ID | P-Value |
| --- | --- | --- |
| p53 signaling pathway | dre04115 | 0.00174693107042 |
| Apoptosis | dre0421 | 0.00204127662297 |
| Necroptosis | dre04217 | 0.00204127662297 |
| Herpes simplex virus 1 infection | dre05168 | 0.00204127662297 |
| Cytokine-cytokine receptor interaction | dre04060 | 0.00204127662297 |
| MAPK signaling pathway | dre04010 | 0.00599480745402 |

### Phylogeny based on orthologous genes

We filtered the clusters that comprised a single sequence from each of the 14 transcriptomes and retrieved 1,369 putative single-copy orthologous genes. We concatenated and aligned these genes into a supermatrix with 1,471,983 informative sites for the 14 taxa. The maximum likelihood tree inferred for each orthologous gene revealed that they formed five well-supported clades based on bootstrap support values (Figure 4). Three notable discrepancies with previous phylogenies were the positions of early diverging *Sinocyclocheilus* lineages: *S. xunlensis*, *S. oxycephalus*, and *S. furcodorsalis*. Our phylogenetic tree unambiguously showed that *S. oxycephalus* was a separate lineage, while *S. xunlensis* was recovered as the sister group of *S. guilinensis*; four species including *S. furcodorsalis* constituted a separate lineage.

**Figure 4.**
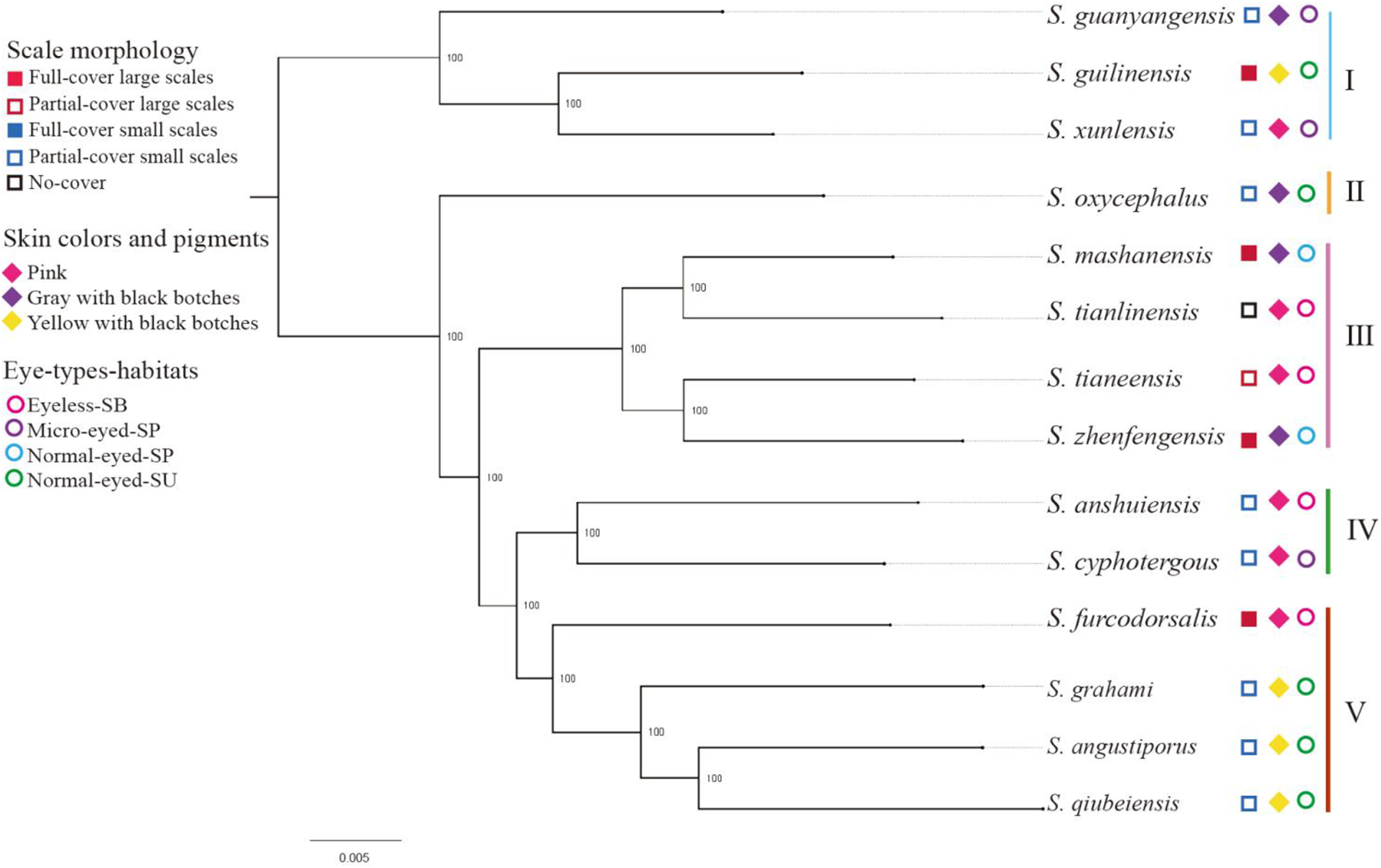
Phylogeny of 14 *Sinocyclocheilus* species with the main clades and Lineages designated by I-V. The maximum likelihood tree was derived from the series data of 1,369 single-copy orthologous genes; bootstrap support is displayed at nodes.

### The skin color

The analysis of color variation in the 14 *Sinocyclocheilus* species highlighted a rich diversity in body coloration and patterns, with each species exhibiting unique colors, mainly in a combination of pinkish-white, gray, and yellow (Figure 5, Supplementary Figure S2).

**Figure 5.**
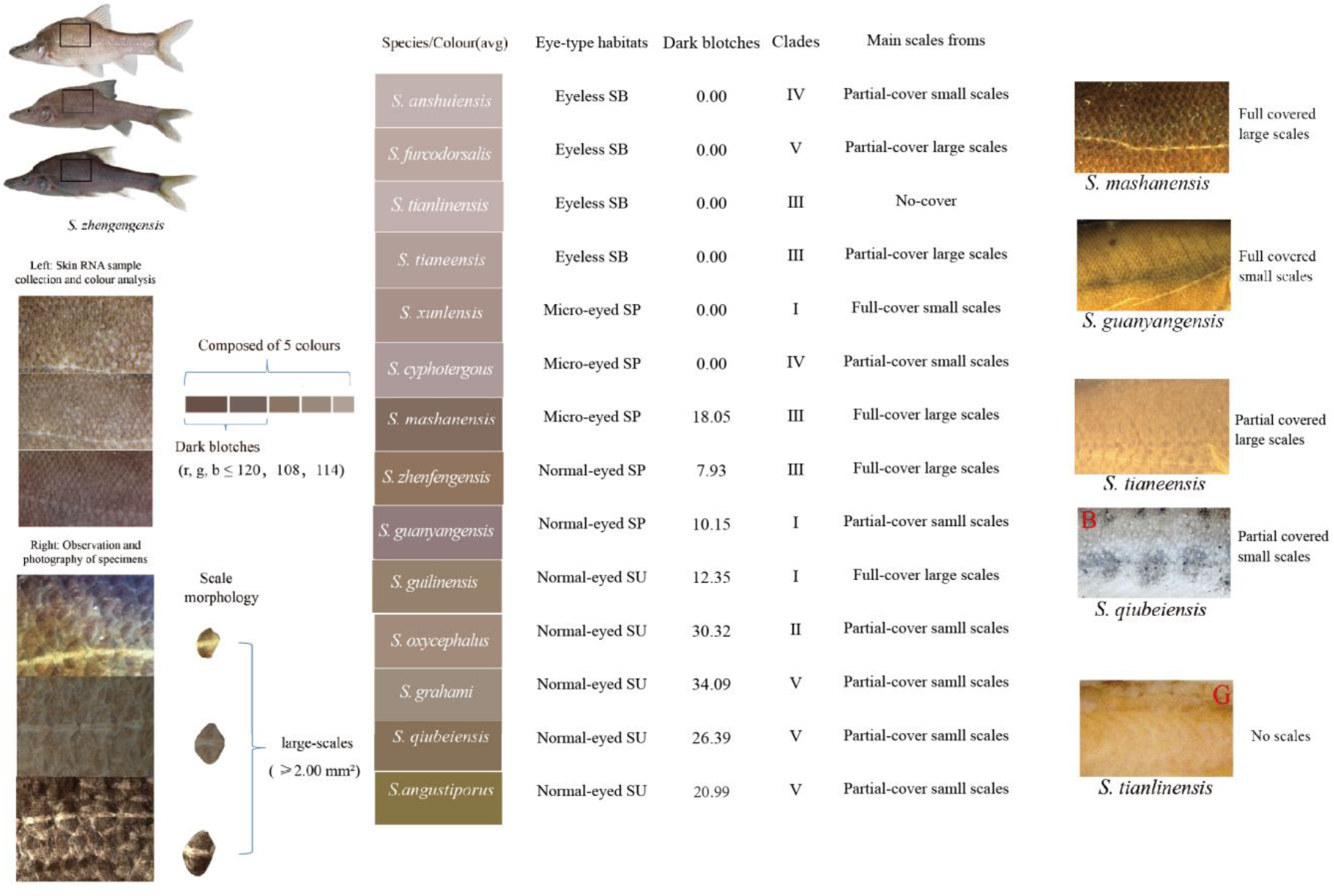
Color and scales characteristics of the 14 *Sinocyclocheilus* species.

For every species, color attributes were assessed based on average skin color value and the proportion of dark blotches and distinct spots on the skin. *Sinocyclocheilus tianlinensis* had the highest skin RGB value (195, 181, 177), followed by *S. anshuiensis* (191, 179, 177) and *S. furcodorsalis* (188, 163, 155); *S. grahami* (138, 123, 111), *S. oxycephalus* (142, 116, 90), and *S. qiubeiensis* (142, 127, 104) displayed the lowest RGB values (Supplementary Table S9). The RGB values for black blotches in these species were under (120, 108, 114) with lower values signifying darker colors, such as dark blotches and spots (Figure 5, Supplementary Table S9).

Skin colors were associated with eye-types and habitats. Normal-eyed SUs had yellow or gray skin, while most SPs exhibited gray or pale gray tones. Unique spots were also indicative of eye-type and habitat groups; Normal-eyed *S. grahami* (SU) had the highest proportion of black spots, succeeded by Normal-eyed *S. oxycephalus* (SU) and Normal-eyed *S. qiubeiensis* (SU). Eyeless SBs and Micro-eyed *S. xunlensis* (SP), which displayed depigmented pink-white skin, had the lowest percentage of dark blotches (Figure 5, Supplementary Table S9). However, microscopic observations showed that SBs and *S. xunlensis* (SP) still had numerous tiny black blotches dispersed across their skin (Supplementary Figure S2); comparatively, Eyeless SBs were pink with fewer pronounced dark blotches and distinct spots.

When examining color in the context of phylogeny, Clades Ⅰ-Ⅴ displayed various combinations of skin color and pigmentation relative to eye-types and habitats (Figure 5, Supplementary Figure S2). Notably, in Clade Ⅲ and Clade Ⅳ, which contain the largest proportion of exclusive cave dwellers, SBs were pink with fewer dark blotches. Hence, the *Sinocyclocheilus* genus adapted to cave environments and evolved with convergent skin coloration in different evolutionary clades.

### Scale characteristics

Microscopic examination of scale characteristics showed a marked reduction in scale size and coverage in 11 *Sinocyclocheilus* species except for two species (*S. mashanensis*: mean ± SD: 3.74 ± 1.36*, S. zhenfengensis*: mean ± SD: 2.93 ± 0.95), which exhibited complete coverage by large scales, and the mean number of their lateral line scales are respectively 49 ± 3 and 43 ± 4. *S. oxycephalus* (mean ± SD: 0.17 ± 0.11), *S. qiubeiensis* (mean ± SD: 0.61 ± 0.28), and *S. grahami* (mean ± SD: 0.72 ± 0.40) had smaller scales, also with more lateral line scales (mean ± SD: 70 ± 3, 78 ± 3 and 72 ± 2), but most of the body scales were buried in the skin and disappeared (Figure 5, Supplementary Figure S2). We found that larger scales and fewer lateral line scales were mainly present in Clade Ⅲ, Ⅳ while smaller scales and more lateral line scales species were mainly found in Clade Ⅴ. The reduction of scale coverage can be found in all clades (I-Ⅴ). This may indicate convergent evolution of scale reduction in *Sinocyclocheilus*.

Generally, SBs exhibited smaller and fewer scales than SPs. The scales of *S. tianlinensis* (SB) were absent altogether, and *S. anshuiensis* (SB) had only rare scales. These species in Clade Ⅲ, Clade Ⅳ revealed a pattern of scales, characterized by a gradual reduction in scale size, decreased coverage and the number of lateral line scales, and eventual disappearance, which may be related to cave adaptation. Clade Ⅴ species displayed variability in scale morphology and coverage, suggesting a chaotic pattern of scale evolution in Clade Ⅴ unrelated to habitat. Here, the small scales were widely spaced (Figure 5).

### Gene expression patterns and skin morphology

The PCA analysis and cluster analysis of the 14 *Sinocyclocheilus* species using expression data from 7,374 orthologous genes revealed that 13 species clustered together, separate from *S. angustiporus* (Figure 6A, Supplementary Table S10). In addition, Micro-eyed SPs and Eyeless SBs showed differences in gene expression compared with Normal-eyed SUs (Figure 6B, Supplementary Table S10). This suggested that patterns of gene expression were similar between these species, but the effects of habitats can still be seen.

**Figure 6.**
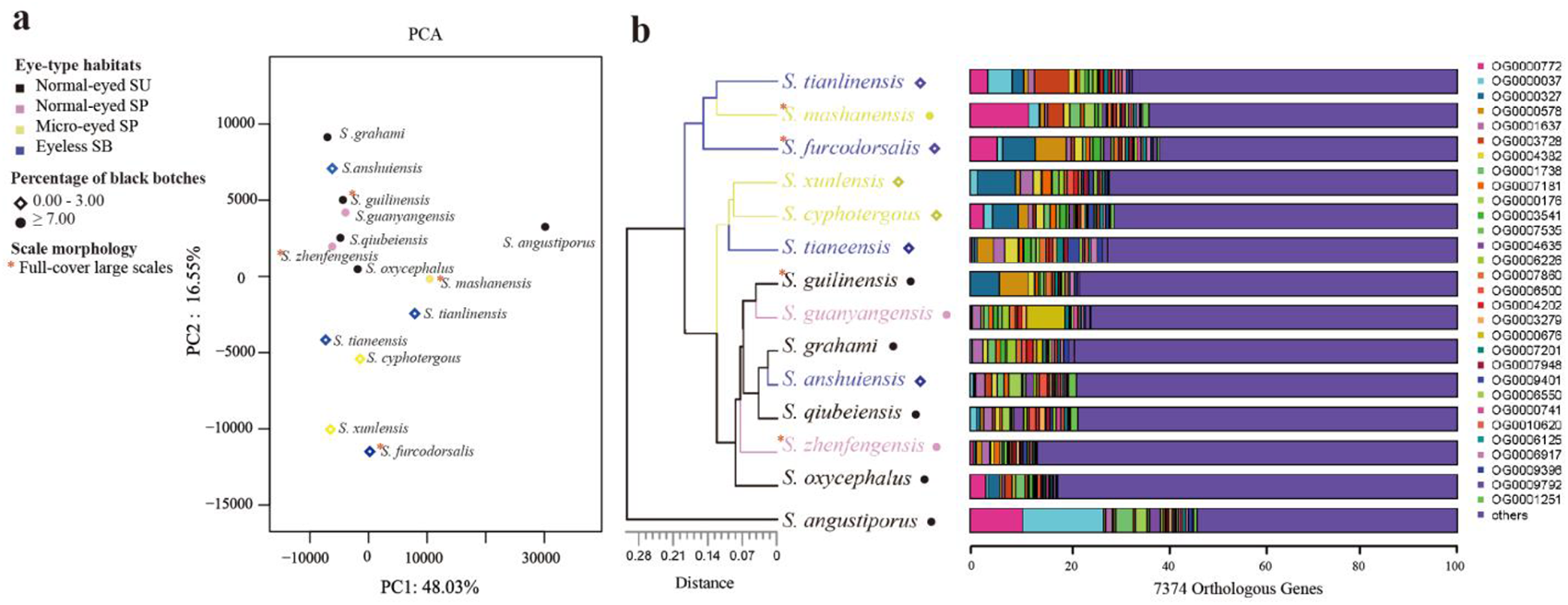
Gene expression clustering of the 14 *Sinocyclocheilus* species based on 7374 orthologous genes. (A) PCA plot showing the relationship between gene expression patterns and habitats/skin morphologies. Together, PC1 and PC2 explain 64.58% of the variability. (B) The stacked bar diagram of cluster tree showing the expression differences of the top 30 orthologous genes with expression changes in different species, and the relationship between these orthologous gene expression changes and habitats/skin morphology.

Interestingly, the second principal component (variance explained 16.55%) differentiated species with and without black blotches (Figure 6A). Moreover, *S. tianeensis*, *S. cyphotergous* and *S. xunlensis* with little black blotches were clustered in one branch, but expression patterns vary among species with full-cover large scales (Figure 6B). This suggested that changes in gene expression patterns may affect pigmentation similarly but not scales.

## DISCUSSION

Broadly, we investigated the skin-related morphology (color and scale characteristics), gene expression patterns, and functional enrichment of DEGs and PSGs among various morphotypes of *Sinocyclocheilus* species, representative of the phylogeny, living in three habitat types (SU, SP, and SB). Our results suggested that habitats may influence changes in color and scale characteristics, gene expression and their function, and hence a driver of skin evolution. Here we discuss our findings and their implications for understanding the broad scale patterns of adaptation of *Sinocyclocheilus* skin for cave-dwelling.

### Possible adaptive mechanisms to different habitats

In our analysis of DEGs, we identified distinct patterns pertaining to metabolism, oxidative stress, and immune responses in various *Sinocyclocheilus* species. These patterns appear to be correlated with their specific habitats. Such results substantiate the presence of cave-environmental gradients within natural ecosystems. In line with our findings, several studies have pointed toward physiological and metabolic distinctions between surface dwelling and cave dwelling species (Krishnan et al., 2020, Stahl and Gross, 2017, Boggs and Gross, 2021, Medley et al., 2022, Yang et al., 2016).

These variances could be due to differences in habitat characteristics like light exposure, ambient oxygen levels, and nutrient resource availability(Garcia-Reyero et al., 2012, Passow et al., 2017, Riddle et al., 2018). However, it is important to note that environmental influences on gene expression are multifaceted and do not merely accumulate in a linear fashion. The interaction among these factors further amplifies the complexity of environmental gradients, making prediction of gene expression responses more challenging (Garcia-Reyero et al., 2012, Passow et al., 2017, Riddle et al., 2018).

While the analysis of coloration and large-scale gene expression patterns may provide some insights, they fall short of identifying the specific environmental factors driving convergent evolution in these habitats. Instead, a more promising approach might involve pinpointing and investigating DEGs, paired with functional annotations informed by a priori hypotheses. This strategy could offer deeper insights into the range of environmental stressors within a habitat, and how they might shape the evolution of different phenotypes.

### Metabolic differences among species across habitats

A large number of GO terms related to energy metabolism (e.g., lipid metabolism, fatty acid metabolism, carbohydrate metabolism, mitochondrial respiration, etc.) were enriched in the comparison of species from different habitats, which were further supported by common KEGG enrichment pathways, such as energy-related Carbon metabolism, Pentose phosphate pathway, PPAR signaling pathway, Fatty acid metabolism, Fatty acid degradation, and OXPHOS.

Most of these DEGs in energy metabolism-related pathways were up-regulated in SUs. This may indicate that SUs, that live in resource rich environments compared to cave dwellers, have a higher energy metabolic rates. In fact, the lowering of metabolism is a well-known feature of organisms living in resource poor cave-environments (Soares and Niemiller, 2020). For instance, *Astyanax* cave-morphs have a lower oxygen consumption and metabolic rate compared to their surface morphs (Boggs and Gross, 2021, Moran et al., 2014). Many of these genes were associated with glycolysis such as fructose-1, 6-diphosphatase (OG0001025), glyceraldehyde-3-phosphate dehydrogenase (OG0001738), 6-phosphogluconate dehydrogenase (OG0011314) and glucose-6-phosphate 1-dehydrogenase (OG0011750) (Okar and Lange, 1999, Randhawa et al., 2014). One of the reasons for this may be due to an abundance of UV light in surface habitats stimulating the skins of SUs leading to enhanced glycolysis (Randhawa et al., 2014).

The differences between cave dwellers (SP and SB) were mainly due to OXPHOS and fatty acid degradation related genes, where they were upregulated in SPs. In addition, under the condition of using fat as an energy source, the expression of the pyruvate dehydrogenase complex-related genes (OG0011863) was upregulated; this is known to contribute to the dynamic balance of glycolysis and tricarboxylic acid cycles (Gray et al., 2014, Pham et al., 2022). The enhanced mitochondrial activity in the skin of cave fish may reflect an increased allocation to detect the environment using non-visual sensory organs, such as lateral line organs, neuromasts and other detectors in the skin, which are enhanced in some cavefish, including in *Sinocyclocheilus* (Yoshizawa et al., 2010; Chen et al., 2022).

In contrast, SBs mainly have enhanced glycolytic processes, as well as enhanced expression of enzymes and proteins related to lipid synthesis and transport, such as *fatty acid synthase* (OG0001380), *very long chain fatty acid elongation protein 6* (OG0010037), *fatty acid-binding protein* (OG0003744). We also found that troglomorphic traits, such as changes in lipid and energy metabolism, appeared to be linked to increased carbohydrate and fat synthesis processes in SPs and SBs, which promotes fat storage (Lam et al., 2022, Xiong, 2021). Overall, the reduced metabolism of nutrients and mitochondrial respiration in SBs may help *Sinocyclocheilus* lower their energy consumption, and increase energy storage, facilitating adaptation to a resource-depleted cave environment (Riddle et al., 2018).

### Immune responses in species from different habitats

Fish skin is acutely sensitive to alterations in the aquatic environment, and it is noteworthy that differentially expressed genes (DEGs) identified within the skin are also responsive to environmental stimuli. Results of the enrichment analysis showed a wide variation in immune mechanisms but with general enrichment in biological processes pertinent to stimulus response, leukocyte proliferation, and apoptosis. Our results also agrees that fish skin has an immune function and highlight the critical role of macrophages during infection (Bangert et al., 2011). In addition to the significant co-enrichment of phagocytosis and inflammatory regulation pathways, we also found that species in different habitats had different immune responses, which may be related to the greater contribution of microbial stimulation and oxygen concentration. Enhanced cave microbial diversity and stimulation in surface water environments may have shaped stronger adaptive immunity in SUs (Rook et al., 2003). In contrast, nutrient limitation and reduced dissolved oxygen in cave water environments may contribute to increased susceptibility to pathogens and risk of inflammation to cave dwellers (SP and SB) (McCormick and Larson, 2008, Taylor and Colgan, 2017).

Compared with cave dwellers (SP, SB), more pro-inflammatory factors such as tumor necrosis factor, C-X-C chemokine, and complement factor related genes were up-regulated in SUs in immune-related pathways. Moreover, genes that were significantly upregulated in microbial immune-related pathways included: *ATPase genes* (*V-type proton ATPase 116 kDa subunit a*: OG0001307; *V-type proton ATPase subunit B*: OG0009257), which were involved in lysosomal function and autophagy flux. The expression of *the vesicle-trafficking protein SEC22b-B* (OG0002841) was also upregulated. These results may indicate an enhanced ability of macrophages to deliver in the skins of SUs, and, along with T cells, support the function of adaptive immunity. In addition, the vesicles were subsequently internalized by macrophages, playing a key role in inflammation resolution. Generally, the macrophages and adaptive immunity in the skins of SUs contributed to microbial resistance (Mohanty and Sahoo, 2010, Lü et al., 2012). The reason for this difference was that there were significant differences in microbial metabolism between different habitats in *Sinocyclocheilus.* we observed upregulation of *inositol-3-phosphate synthase 1-A-like isoenzyme X2* (OG0003974) and *UDP-N-acetylglucosamine pyrophosphorylase* (*UAP1*, OG0008682), indicating increased eukaryotic growth and reproductive activity in skin microorganisms (Reynolds, 2009, Behr, 2011). The dynamics and intensity of viral replication may tend to weaken due to the lowering of temperature within the cave environment (Demory et al., 2017). In fact, it is generally known that caves are a biodiversity-depleted environment, including that of pathogens (Peuß et al., 2020). Thus, investment strategies for microbial immunity in cave-dwelling species may be lower compared to those of SUs.

There is a disparity between cave-dwelling species (SP, SB) with respect to the expression levels of genes involved in hypoxia (ECM-receptor interaction pathway and hematopoietic cell lines pathway) and inflammatory (complement and coagulation cascades pathway) related responses and pathways (Chen et al., 2021, Morikawa and Takubo, 2016). Inflammation plays an important role in the immune response, serving as a critical pathophysiological reaction of the organism to pathogenic invasion, tissue damage, and other stimuli. Hypoxia also emerges as a significant modulator of both inflammatory and immune responses (Zhao et al., 2016, Bhatti et al., 2017, Duan et al., 2022). In contrast to SPs, a large number of proinflammatory factors (such as complement factors, coagulation factors, and integrin) were up-regulated in SBs. This suggested that SBs had a stronger innate immune response to microbes. Interestingly, we found the least immune-associated GO between SBs and SPs, especially T-cellular immune-related. Thus, cavefish may reduce investment in immune cells, but enhance the sensitivity of the innate immune system as suggested by some previous studies as well (Mayer et al., 2016, Peuß et al., 2020). Hence, our findings suggest that SBs may employ adaptive strategies in cave environments that involve a reduction in T-cell mediated immune responses, but an enhancement in the regulation of innate immune defenses. Moreover, efficient resolution of inflammation and infection in the dermal tissues of SBs necessitates spontaneous apoptosis, induced by microenvironmental factors, to facilitate the clearance of abundant neutrophils at the site of infection.

### Oxidative stress in species from different habitats

Though both inflammatory signaling and the mitochondrial electron transport chain (ETC) contribute to the generation of cellular reactive oxygen species (ROS), the mitochondria principally serve as the primary site of cellular ROS production. Additionally, they are the primary targets of a multitude of exogenous toxic effects stemming from environmental chemical agents and ROS themselves (Li et al., 2017). In our study, cave dwellers (SP, SB) were significantly enriched for most of the diverse antioxidant stress-related pathways. The lower dissolved oxygen concentrations and nutrients in the cave water environment, as demonstrated in previous studies, constitute a significant limiting factor (Boggs and Gross, 2021, Riddle et al., 2018). This could contribute to the oxidative stress observed in cave-dwelling species (SP, SB). Examination of genes in the anti-oxidative stress-related pathways of SUs and SBs revealed slight differences in expressions between them. Some of the minor differences were that SUs mainly upregulated genes in the Peroxisome pathway, while SBs mainly enhanced genes in the P450 and Glutathione metabolism pathway to enhance cellular antioxidant action and integrated detoxification. The increased activity of catalase in SUs contributes to mitochondrial respiration and enhances fatty acid oxidation and utilization (Yu et al., 2003). And cytochrome P450 may target SBs in their sensitivity to exogenous chemicals (including pesticides) in the cave environment, as well as the need for thermoregulation (Guengerich et al., 2016). In a previous study, cavefish showed increased levels of stress compared to surface fish, and the expression of genes involved in glutathione metabolism was also increased to prevent oxidative stress under prolonged fasting (Krishnan et al., 2020, Medley et al., 2022).

Interestingly, almost all oxidative stress-related genes in the Peroxisome pathway and P450 pathway were down-regulated in SP skins, suggesting that they may have a stronger environmental oxidative stress resistance. It has been show that the down-regulation of stress-related genes in response to abiotic or biological changes in the environment is an indicator of stress-resistant populations and species (Bailey et al., 2017). Previous studies had shown the heterogeneity of SP environments may enhance their immune capacity (Yang et al., 2016). The extent to which environmental stressors influence appears to be lesser and is contingent upon the physiological strategies adopted by the species; furthermore, microevolutionary processes contribute to the augmentation of resistance of an organism (Sun et al., 2015).

SPs reduced tissue hypoxia by enhancing oxidative phosphorylation (OXPHOS), thereby enhancing mitochondrial respiration and ATP synthesis, thereby enhancing adaptation to hypoxia. Cytochrome c oxidase (COX) was a key enzyme in establishing a more efficient mitochondrial respiratory chain (MRC) to improve oxygen utilization under hypoxic conditions. This may help SPs maintain homeostasis of mitochondrial regulation of energy production, reactive oxygen species homeostasis, and cell death in the hypoxic microenvironment caused by skin immunization (Garvin et al., 2015, Heather et al., 2012). Our study demonstrated different hypoxic adaptation strategies in cave dwellers, implicating that alterations in mitochondrial respiration rate and enzymatic activity might play pivotal roles in the regulation and maintenance of redox homeostasis.

### Transcriptional Plasticity and Convergent Evolution

Our ML tree, based on 1,369 unique single-copy orthologous genes, showed some discordance when compared to recent mitochondrial gene (mt-DNA) and RAD-seq based studies (Zhao and Zhang, 2009, Wen et al., 2022, Xu et al., 2023, Mao et al., 2021, Mao et al., 2022b). The robust node support in our phylogeny might be attributed to the abundance of genes under selection derived from transcriptome sequencing. However, this precision may have a downside: the unique genes associated with the transcriptome we used could potentially not be under neutral evolution, thereby causing the phylogeny of expressed genes important for the functions of the skin. Most notably, Clade I of our study contains species from three clades that employed mt-DNA/RAD-seq data. This was possibly due to shared skin traits. In addition, *S. oxycephalus* formed a distinct divergent lineage (II). The skin of these species had a distinctive gray color. And previous studies have shown that its skin is charcoal gray skin unlike other types (Li et al., 2020). Furthermore, we identified *S. furcodorsalis*, which was from clade B of previous studies placed in our Clade V, which contained predominantly species from clade D (which are mostly SUs) (Mao et al., 2021). This suggested that *S. furcodorsalis* may have a skin transcription profile similar to that of SUs. In fact, this species, despite being a cave-dweller, enigmatically had some of the most prominent scales. However, our phylogeny lacked high taxon sampling, which prevented us from making deeper inferences on transcriptional plasticity. However, this study highlights the significance and precision of transcriptomic data in understanding the evolutionary relationships among *Sinocyclocheilus* species.

It is known that the phenotypic adaptations of cave-dwelling in *Sinocyclocheilus* cavefish involve changes in eye types and color and pigmentation patterns (Li et al., 2020, Meng et al., 2013); our results from the context of skin morphology and gene expression confirm this. We found that Eyeless SBs and some Micro-eyed SPs without obvious black blotches showing similar expression patterns, even though they were in different clades (Figure 5). Existing literature has established that changes in skin color, driven primarily by adaptation to diverse habitats, serve as a major contributing factor to differential gene expression patterns (Li et al., 2020, Hinaux et al., 2013, Stahl and Gross, 2015, Gross et al., 2016). Skin color adaptations in fish are known to occur rapidly when they are introduced to novel environments (Leclercq et al., 2010, Nilsson Sköld et al., 2013).

The relationship between fish skin color and melanin content was examined, focusing on key genes involved in phenylalanine, tyrosine and tryptophan biosynthesis pathway, and especially the tyrosine metabolism (Figure 7). We found that the tyrosine metabolic pathway, influenced by habitat, played a role in determining melanosis in the skin of *Sinocyclocheilus*. Previous research has demonstrated that decreased *phenylalanine-4-hydroxylase* (*PAH,* OG: 0000390) activity can affect *Try* activity (Leandro et al., 2017), and inhibition of phenylalanine conversion to tyrosine will lead to reduced melanin production (Staudigl et al., 2011, Zhou et al., 2021). However, we found low expression of the gene encoding *PAH* in these species. This may be due to the lack of phenylalanine and protein catabolic inputs in the diet of cavefish (Borowsky, 2018). However, L-phenylalanine (L-Phe) is also an essential protein-producing amino acid, and it is important to avoid it being completely catabolic (Xiao et al., 2020). Therefore, in order to achieve the dual role of effective preservation and removal of excess L-Phe, this regulatory mechanism may be more favorable to the adaptation of their environment.

**Figure 7.**
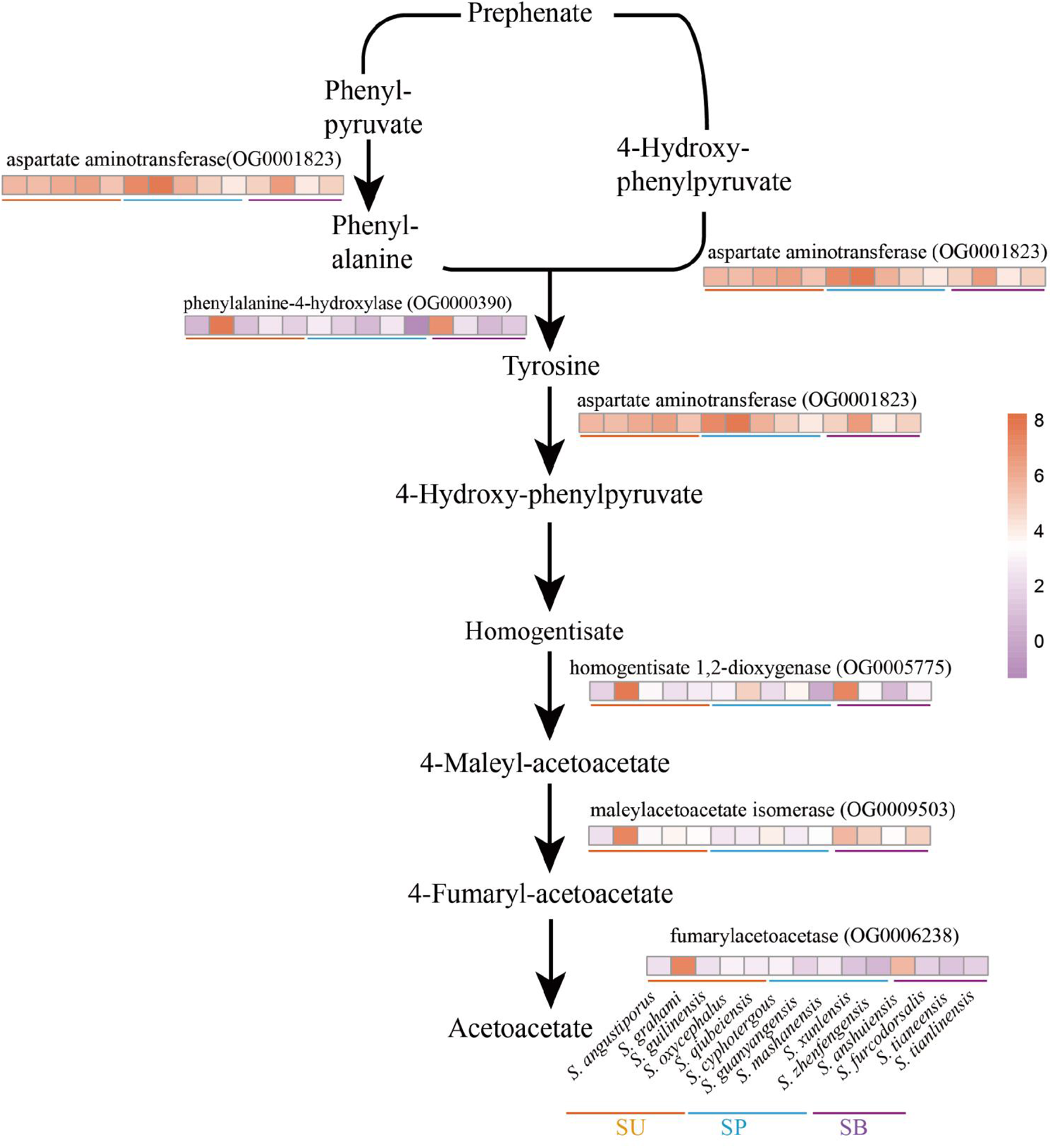
Flow of phenylalanine metabolism and tyrosinase metabolism and heat map of related enzyme gene expression in 14 *Sinocyclocheilus* species. The darker the orange, higher the expression; the darker the purple, lower the expression.

Our analysis revealed that SBs had the fewest black blotches and the lowest expression of tyrosine metabolism-related enzymes: *aspartate aminotransferase* (*AST*, OG0001823), *fumarylacetoacetase* (*FAH*, OG0006238; Figure 5, 7). This indicated that phenylalanine and tyrosine metabolism was inhibited, and the final melanin synthesis was affected. Consistent with previous research, melanin synthesis is triggered by sunlight, which can offer benefits such as camouflage, protection from ultraviolet radiation, and a role in social signaling (D’Mello et al., 2016, Rzepka et al., 2016). Thus, whereas melanin synthesis is inhibited in SBs that inhabit lightless environments.

SPs such as *S. guanyangensis* and *S. cyphotergous* had elevated *AST* activity. The way these species produce melanin may be related to the accumulation of homogentisic acid (HGA). HGA oxidation leads to melanin-like pigmentation (Giustarini et al., 2012). Interestingly, cultivation of these species in our laboratory revealed that SPs seemed to exhibit an enhanced ability to adapt rapidly to fluctuating light conditions. This may enable them to adapt to the changing light environment between cave and semi-enclosed cave environments. Therefore, *AST* may have an important role in fish skin differentiation and variation through the melanogenesis pathway. The observed pigmentation differences among *Sinocyclocheilus* species may represent adaptations to their specific environments. Reduced pigmentation could be advantageous in the dark cave environment, while increased pigmentation may be more beneficial for survival in semi-enclosed and surface environments (Romero and Green, 2005, Soares and Niemiller, 2020).

Degeneration of scales was found in all clades in our study (Figure 4). It is not unexpected to find skin scale degradation prevalent in *Sinocyclocheilus*, as it is a trait seen in some other Carpiformes, unrelated to cave environments (Zhao and Zhang, 2009, Zhu et al., 2019, Harris et al., 2008). The scales of *Sinocyclocheilus* are known to be bony scales, and the posterior part of their scales consists of bone (Zhao and Zhang, 2009). We found that *Reticulocalbin 3, EF-hand calcium binding domain* (OG0016528)*, S100 calcium binding protein U* (OG0016522), and *Positive regulation of vitamin D receptor signaling pathway* (OG0016683) were under positive selection, which may contribute to the calcium-bone homeostasis of fish skin (Schäfer and Heizmann, 1996, Girard et al., 2015, Bouillon and Suda, 2014). And PSGs are enriched in the MAPK signaling pathway, which is known to be associated with scale development, so the role of such genes involved in scale evolution is strong (Zhou et al., 2022a); there is also a strong association between scale regression and cave-dwelling. However, a deeper analysis is warranted to explore the connection between cave environments and scale features perse, i.e. a comparison between scaleless *Sinocyclocheilus* and scaleless other Carpiformes. Our research showcases the remarkable adaptability of cave-dwelling species to dynamic environments, offering insights into fish adaptation processes in response to anthropogenic alterations, such as habitat degradation and climate change. *Sinocyclocheilus* cavefish share few positive selections among their skin orthologous genes, and genes associated with the limited skin phenotype are more difficult to detect. The most likely plausible reason for this evolutionary phenomenon is the low rate of evolution of the cavefish genome and genome-wide relaxation of selection(Policarpo et al., 2020, Torres-Paz et al., 2018, Zhao et al., 2022). Our study demonstrates the remarkable adaptability of cave-dwelling species to dynamic environments and provides insights into the process of fish adaptation to anthropogenic changes such as habitat degradation and climate change.

PSGs in *Sinocyclocheilus* were enriched in herpes simplex virus 1 (HSV-1) infection, cytokine-cytokine receptor interactions, p53 signaling pathway, necrosis, and apoptosis related pathways. The current study has found that under prolonged environmental stress (e.g. hypoxia, prolonged hypothermia, starvation, etc.), facilitation and evolution of pathways such as p53 signaling pathway, MAPK signaling pathway, cytokine-cytokine receptor interaction, and apoptosis related pathways are prone to occur and echoes our study (Voskarides et al., 2022, Mao et al., 2022a, Tong et al., 2021, Li et al., 2023). Our study highlighted that an important aspect of skin evolution is resistance to viral infection. These pathways promote coordination between cells and contribute to the clearance of pathogenic infections. Previous studies have shown that p53 plays a dual role in the replication of HSV-1 at different stages of infection (Voskarides et al., 2022, Sato and Tsurumi, 2013). Our study found that one of the p53 homologs: the *Tumor protein p63 regulated 1* (OG0018161) was subject to the strongest positive selection (Aloni-Grinstein et al., 2018). Interestingly, it has been shown that HSV-1 manages to counteract this negative effect of p53 through the viral protein ICP22, which is able to bind p53 directly and eliminate its function (Sato and Tsurumi, 2013). Like many other viruses, herpesviruses their hosts co-evolved (Chawla et al., 2022). Fish skin is the first line of defense against viral and pathogenic microbial infections in the aquatic environment, and the adaptive evolution of immune and apoptosis-related genes may have facilitated the development of immune function in the skin of *Sinocyclocheilus*, enabling their ability to rapidly adapt to new environments and to rapidly occupy vacant ecological niches.

## CONCLUSIONS

In conclusion, our study focused on investigating the genetic basis of environmental adaptation in *Sinocyclocheilus* cavefish, specifically exploring the influence of gene regulation on skin-related traits. Through a radiation-scale analysis involving representatives from major clades and different habitat types, we uncovered important findings regarding the adaptive mechanisms of these cave-dwelling fish.

We observed that different habitats exerted significant influences on color and scale characteristics, gene expression patterns, and functional enrichment of differentially expressed genes (DEGs) and positively selected genes (PSGs) in *Sinocyclocheilus* species. The functional enrichment analysis revealed distinct patterns in signaling mechanisms, oxidative stress, energy metabolism, and immune responses associated with different habitats.

Regarding metabolic differences among species across habitats, we found that species from surface habitats (SUs) exhibited higher energy metabolic rates, reflected in enriched energy metabolism pathways such as carbohydrate metabolism, amino acid metabolism and energy meta signaling mechanisms.

Conversely, SPs enhanced fat decomposition and mitochondrial respiration, while SBs showed enhanced glycolysis processes, lipid synthesis, and transport pathways, possibly as an adaptation to energy-poor cave environments.

The immune responses of species from different habitats also exhibited variations. SUs displayed the mechanism by which macrophages combine to adapt immune resistance to microbial infection, indicating a stronger response to microbial challenges. In contrast, cave dwellers showed differences in immune response patterns, with SBs exhibiting stronger innate immune responses to microbes. Additionally, oxidative stress-related pathways were enriched in cave dwellers, potentially as a response to the unique challenges of their low-oxygen cave environments.

The analysis of transcriptional plasticity and convergent evolution in *Sinocyclocheilus* cavefish revealed interesting findings. Skin color adaptations were associated with differential gene expression patterns, particularly in the tyrosine metabolism pathway, which influences melanin production. The presence or absence of scales in different species was also observed, with scale regression prevalent in cave-dwelling species. These observations highlight the significance of skin morphology and gene expression in understanding the evolutionary relationships among *Sinocyclocheilus* species.

This research demonstrates the adaptability of cave-dwelling species to dynamic environments. The knowledge generated maybe useful also in predicting the processes of fish adaptation in response to habitat degradation and climate change. The genetic mechanisms underlying environmental adaptations, such as metabolic adjustments, immune responses, oxidative stress regulation, and phenotypic traits, contribute to our understanding of the broad-scale patterns of adaptation in *Sinocyclocheilus* skin for cave-dwelling. Further investigations are warranted to deepen our knowledge of skin evolution and its adaptive significance in this unique group of cavefish.

## Author Contributions

XL and MM: Conceptualization. XL, TM, YL: Data curation. XL, BC, TM, YL and MM: Methodology. XL: Software, formal analysis, data curation, visualization. XL, BC and MM: Writing the original draft. MM: Project administration and supervision. JY and MM: Resources. XL, MM, CB, TM, JY and YL: writing—review and editing. All authors contributed to the article and approved the submitted version.

## Ethics statement

The animal study was reviewed and approved by Institutional Animal Care and Use Committee of Guangxi University (GXU), Nanning-China (#GXU2021-126).

## Acknowledgments

We thank the following individuals: Juntao Hu (Fudan University) and Jian Yang for suggestions to improve the paper. Madhava Meegaskumbura and Bing Chen for helpful comments on an earlier version of the manuscript; Tingru Mao for technical assistance; Yewei Liu, Shipeng Zhou and Dan Sun for assistance in the field.

## REFERENCES

Abràmoff, M. D., Magalhães, P. J., Ram, S. J. (2004). Image processing with ImageJ. Biophotonics international, 11, 36–42. doi: 10.3233/ISU-1991-115-601

Aloni-Grinstein, R., Charni-Natan, M., Solomon, H., Rotter, V. (2018). p53 and the Viral Connection: Back into the Future (‡). Cancers (Basel), 10. doi: 10.3390/cancers10060178

Andrews, S. (2010). FastQC: A quality control tool for high throughput sequence data. Available online at: https://www.bioinformatics.babraham.ac.uk/projects/fastqc/.

Ángeles Esteban, M. (2012). An overview of the immunological defenses in fish skin. International scholarly research notices, 2012. doi: 10.5402/2012/853470

Austin, B. (2006). The bacterial microflora of fish, revised. TheScientificWorldJOURNAL, 6, 931–945. doi: 10.1100/tsw.2006.181

Bailey, A., De Wit, P., Thor, P., Browman, H. I., Bjelland, R., Shema, S., Fields, D. M., Runge, J. A., Thompson, C., Hop, H. (2017). Regulation of gene expression is associated with tolerance of the Arctic copepod Calanus glacialis to CO_2_-acidified sea water. Ecology and Evolution, 7, 7145–7160. doi: 10.1002/ece3.3063

Bangert, C., Brunner, P. M., Stingl, G. (2011). Immune functions of the skin. Clinics in dermatology, 29, 360–376. doi: 10.1016/j.clindermatol.2011.01.006

Behr, J.B. (2011). Chitin synthase, a fungal glycosyltransferase that is a valuable antifungal target. Chimia, 65, 49–49. doi:10.2533/chimia.2011.49

Belkaid, Y., Hand, T. W. (2014). Role of the microbiota in immunity and inflammation. Cell, 157, 121–141. doi: 10.1016/j.cell.2014.03.011

Bhatti, J. S., Bhatti, G. K., Reddy, P. H. (2017). Mitochondrial dysfunction and oxidative stress in metabolic disorders—A step towards mitochondria based therapeutic strategies. Biochimica et Biophysica Acta (BBA)-Molecular Basis of Disease, 1863, 1066–1077. doi: 10.1016/j.bbadis.2016.11.010

Boggs, T., Gross, J. (2021). Reduced oxygen as an environmental pressure in the evolution of the blind Mexican cavefish. Diversity, 13, 26. doi: 10.3390/d13010026

Borowsky, R. (2018). Cavefishes. Current Biology, 28, R60–R64. doi: 10.1016/j.cub.2017.12.011

Bouillon, R., Suda, T. (2014). Vitamin D: calcium and bone homeostasis during evolution. Bonekey Rep, 3, 480. doi: 10.1038/bonekey.2013.214

Castresana, J. (2000). Selection of conserved blocks from multiple alignments for their use in phylogenetic analysis. Molecular Biology and Evolution, 17, 540–552. doi: 10.1093/oxfordjournals.molbev.a026334

Chawla, K., Subramanian, G., Rahman, T., Fan, S., Chakravarty, S., Gujja, S., Demchak, H., Chakravarti, R., Chattopadhyay, S. (2022). Autophagy in Virus Infection: A Race between Host Immune Response and Viral Antagonism. Immuno, 2, 153–169. doi:org/10.3390/immuno2010012

Chen, B., Mao, T. R., Liu, Y. W., Dai, W. A., Li, X. L., Rajput, A. P., Pie, M. R., Yang, J., Gross, J. B., Meegaskumbura, M. (2022). Sensory evolution in a cavefish radiation: patterns of neuromast distribution and associated behaviour in *Sinocyclocheilus* (Cypriniformes: Cyprinidae). Proceedings of the Royal Society B: Biological Sciences, 289, 20221641. doi: 10.1098/rspb.2022.1641

Chen, G., Pang, M., Yu, X., Wang, J., Tong, J. (2021). Transcriptome sequencing provides insights into the mechanism of hypoxia adaption in bighead carp (Hypophthalmichthys nobilis). Comparative Biochemistry and Physiology Part D: Genomics and Proteomics, 40, 100891. doi: 10.1016/j.cbd.2021.100891

D’mello, S. A., Finlay, G. J., Baguley, B. C., Askarian-Amiri, M. E. (2016). Signaling Pathways in Melanogenesis. International Journal of Molecular Sciences, 17. doi: 10.3390/ijms17071144

Demory, D., Arsenieff, L., Simon, N., Six, C., Rigaut-Jalabert, F., Marie, D., Ge, P., Bigeard, E., Jacquet, S., Sciandra, A. (2017). Temperature is a key factor in Micromonas–virus interactions. The ISME journal, 11, 601–612. doi: 10.1038/ismej.2016.160

Duan, H., Jing, L., Xiang, J., Ju, C., Wu, Z., Liu, J., Ma, X., Chen, X., Liu, Z., Feng, J. (2022). CD146 Associates with Gp130 to Control a Macrophage Pro-inflammatory Program That Regulates the Metabolic Response to Obesity. Advanced Science, 9, 2103719. doi: 10.1002/advs.202103719

Edgar, R. C. (2022). Muscle5: High-accuracy alignment ensembles enable unbiased assessments of sequence homology and phylogeny. Nature Communications, 13, 6968. doi: 10.1038/s41467-022-34630-w

Ellison, A. R., Wilcockson, D., Cable, J. (2021). Circadian dynamics of the teleost skin immune-microbiome interface. Microbiome, 9, 1–18. doi: 10.1186/s40168-021-01160-4

Emms, D. M., Kelly, S. (2015). OrthoFinder: solving fundamental biases in whole genome comparisons dramatically improves orthogroup inference accuracy. Genome Biol, 16, 157. doi: 10.1186/s13059-015-0721-2

Fortune, E. S., Andanar, N., Madhav, M., Jayakumar, R. P., Cowan, N. J., Bichuette, M. E., Soares, D. (2020). Spooky Interaction at a Distance in Cave and Surface Dwelling Electric Fishes. Front Integr Neurosci, 14, 561524. doi: 10.3389/fnint.2020.561524

Fu, L., Niu, B., Zhu, Z., Wu, S., Li, W. (2012). CD-HIT: accelerated for clustering the next-generation sequencing data. Bioinformatics, 28, 3150–3152. doi: 10.1093/bioinformatics/bts565

Garvin, M. R., Thorgaard, G. H., Narum, S. R. (2015). Differential Expression of Genes that Control Respiration Contribute to Thermal Adaptation in Redband Trout (Oncorhynchus mykiss gairdneri). Genome Biology and Evolution, 7, 1404–1414. doi: 10.1093/gbe/evv078

Gibert, J., Deharveng, L. (2002). Subterranean Ecosystems: A Truncated Functional Biodiversity: This article emphasizes the truncated nature of subterranean biodiversity at both the bottom (no primary producers) and the top (very few strict predators) of food webs and discusses the implications of this truncation both from functional and evolutionary perspectives. BioScience, 52, 473–481. doi: 10.1641/0006-3568(2002)052[0473:SEATFB]2.0.CO;2

Girard, F., Venail, J., Schwaller, B., Celio, M. R. (2015). The EF-hand Ca^2+^-binding protein super-family: A genome-wide analysis of gene expression patterns in the adult mouse brain. Neuroscience, 294, 116–155. doi: 10.1016/j.neuroscience.2015.02.018

Giustarini, D., Dalle-Donne, I., Lorenzini, S., Selvi, E., Colombo, G., Milzani, A., Fanti, P., Rossi, R. (2012). Protein thiolation index (PTI) as a biomarker of oxidative stress. Free Radical Biology and Medicine, 53, 907–915. doi: 10.1016/j.freeradbiomed.2012.06.022

Grabherr, M. G., Haas, B. J., Yassour, M., Levin, J. Z., Thompson, D. A., Amit, I., Adiconis, X., Fan, L., Raychowdhury, R., Zeng, Q., Chen, Z., Mauceli, E., Hacohen, N., Gnirke, A., Rhind, N., Di Palma, F., Birren, B. W., Nusbaum, C., Lindblad-Toh, K., Friedman, N., Regev, A. (2011). Full-length transcriptome assembly from RNA-Seq data without a reference genome. Nat Biotechnol, 29, 644–652. doi: 10.1038/nbt.1883

Gray, L. R., Tompkins, S. C., Taylor, E. B. (2014). Regulation of pyruvate metabolism and human disease. Cell Mol Life Sci, 71, 2577–2604. doi: 10.1007/s00018-013-1539-2

Gross, J. B. (2012). The complex origin of *Astyanax* cavefish. BMC evolutionary biology, 12, 1–12. doi: 10.1186/1471-2148-12-105

Gross, J. B., Powers, A. K., Davis, E. M., Kaplan, S. A. (2016). A pleiotropic interaction between vision loss and hypermelanism in *Astyanax mexicanus* cave x surface hybrids. BMC Evol Biol, 16, 145. doi: 10.1186/s12862-016-0716-y

Guengerich, F. P., Waterman, M. R., Egli, M. (2016). Recent Structural Insights into Cytochrome P450 Function. Trends Pharmacol Sci, 37, 625–640. doi: 10.1016/j.tips.2016.05.006

Harris, M. P., Rohner, N., Schwarz, H., Perathoner, S., Konstantinidis, P., Nüsslein-Volhard, C. (2008). *Zebrafish* eda and edar mutants reveal conserved and ancestral roles of ectodysplasin signaling in vertebrates. PLoS genetics, 4, e1000206. doi: 10.1371/journal.pgen.1000206

Heather, L. C., Cole, M. A., Tan, J.J., Ambrose, L. J. A., Pope, S., Abd-Jamil, A. H., Carter, E. E., Dodd, M. S., Yeoh, K. K., Schofield, C. J., Clarke, K. (2012). Metabolic adaptation to chronic hypoxia in cardiac mitochondria. Basic Research in Cardiology, 107, 268. doi: 10.1007/s00395-012-0268-2

Hinaux, H., Poulain, J., Da Silva, C., Noirot, C., Jeffery, W. R., Casane, D., Rétaux, S. (2013). De novo sequencing of Astyanax mexicanus surface fish and Pachón cavefish transcriptomes reveals enrichment of mutations in cavefish putative eye genes. PLoS One, 8, e53553. doi: 10.1371/journal.pone.0053553

Huang, Y., Chain, F. J. J., Panchal, M., Eizaguirre, C., Kalbe, M., Lenz, T. L., Samonte, I. E., Stoll, M., Bornberg-Bauer, E., Reusch, T. B. H. (2016). Transcriptome profiling of immune tissues reveals habitat-specific gene expression between lake and river sticklebacks. Wiley-Blackwell Online Open, 25. doi: 10.1111/mec.13520

Huerta-Cepas, J., Szklarczyk, D., Forslund, K., Cook, H., Heller, D., Walter, M. C., Rattei, T., Mende, D. R., Sunagawa, S., Kuhn, M., Jensen, L. J., Von Mering, C., Bork, P. (2016). eggNOG 4.5: a hierarchical orthology framework with improved functional annotations for eukaryotic, prokaryotic and viral sequences. Nucleic Acids Res, 44, D286–293. doi: 10.1093/nar/gkv1248

Jablonski, N. G., Chaplin, G. (2010). Human skin pigmentation as an adaptation to UV radiation. Proceedings of the National Academy of Sciences, 107, 8962–8968. doi: 10.1073/pnas.0914628107

Jeffery, W. R. (2001). Cavefish as a model system in evolutionary developmental biology. Developmental Biology, 231, 1–12. doi: 10.1006/dbio.2000.0121

Jiang, W. S., Li, J., Lei, X. Z., Wen, Z. R., Han, Y. Z., Yang, J. X., Chang, J. B. (2019). *Sinocyclocheilus sanxiaensis*, a new blind fish from the Three Gorges of Yangtze River provides insights into speciation of Chinese cavefish. Zoological research, 40, 552. doi: 10.24272/j.issn.2095-8137.2019.065

Kanehisa, M., Araki, M., Goto, S., Hattori, M., Hirakawa, M., Itoh, M., Katayama, T., Kawashima, S., Okuda, S., Tokimatsu, T., Yamanishi, Y. (2008). KEGG for linking genomes to life and the environment. Nucleic Acids Res, 36, D480–484. doi: 10.1093/nar/gkm882

Krishnan, J., Persons, J. L., Peuss, R., Hassan, H., Kenzior, A., Xiong, S., Olsen, L., Maldonado, E., Kowalko, J. E., Rohner, N. (2020). Comparative transcriptome analysis of wild and lab populations of Astyanax mexicanus uncovers differential effects of environment and morphotype on gene expression. J Exp Zool B Mol Dev Evol, 334, 530–539. doi: 10.1002/jez.b.22933

Lam, S. M., Li, J., Sun, H., Mao, W., Lu, Z., Zhao, Q., Han, C., Gong, X., Jiang, B., Chua, G. H. (2022). Quantitative lipidomics and spatial MS-Imaging uncovered neurological and systemic lipid metabolic pathways underlying troglomorphic adaptations in cave-dwelling fish. Molecular Biology and Evolution, 39, msac050. doi: 10.1093/molbev/msac050

Langmead, B., Salzberg, S. L. (2012). Fast gapped-read alignment with Bowtie 2. Nat Methods, 9, 357–359. doi: 10.1038/nmeth.1923

Latella, L. (2019). Biodiversity: China. Encyclopedia of caves. (pp. 127–135). Elsevier.

Leandro, J., Stokka, A. J., Teigen, K., Andersen, O. A., Flatmark, T. (2017). Substituting Tyr^138^ in the active site loop of human phenylalanine hydroxylase affects catalysis and substrate activation. FEBS Open Bio, 7, 1026–1036. doi: 10.1002/2211-5463.12243

Leclercq, E., Taylor, J. F., Migaud, H. (2010). Morphological skin colour changes in teleosts. Fish and Fisheries, 11, 159–193. doi: 10.1111/j.1467-2979.2009.00346.x

Li, B., Dewey, C. N. (2011). RSEM: accurate transcript quantification from RNA-Seq data with or without a reference genome. BMC Bioinformatics, 12, 323. doi: 10.1186/1471-2105-12-323

Li, C. Q., Chen, H. Y., Zhao, Y. C., Chen, S. Y., Xiao, H. (2020). Comparative transcriptomics reveals the molecular genetic basis of pigmentation loss in *Sinocyclocheilus* cavefishes. Ecology and Evolution, 10, 14256–14271. doi: 10.1002/ece3.7024

Li, X., Fang, P., Yang, W. Y., Chan, K., Lavallee, M., Xu, K., Gao, T., Wang, H., Yang, X. (2017). Mitochondrial ROS, uncoupled from ATP synthesis, determine endothelial activation for both physiological recruitment of patrolling cells and pathological recruitment of inflammatory cells. Can J Physiol Pharmacol, 95, 247–252. doi: 10.1139/cjpp-2016-0515

Li, X., Wang, X., Yang, C., Lin, L., Yuan, H., Lei, F., Huang, Y. (2023). A de novo assembled genome of the Tibetan Partridge (Perdix hodgsoniae) and its high-altitude adaptation. Integrative Zoology, 18, 225–236. doi: 10.1111/1749-4877.12673

Loomis, C., Peuß, R., Jaggard, J. B., Wang, Y., Mckinney, S. A., Raftopoulos, S. C., Raftopoulos, A., Whu, D., Green, M., Mcgaugh, S. E., Rohner, N., Keene, A. C., Duboue, E. R. (2019). An Adult Brain Atlas Reveals Broad Neuroanatomical Changes in Independently Evolved Populations of Mexican Cavefish. Front Neuroanat, 13, 88. doi: 10.3389/fnana.2019.00088

Lü, A., Hu, X., Xue, J., Zhu, J., Wang, Y., Zhou, G. (2012). Gene expression profiling in the skin of zebrafish infected with Citrobacter freundii. Fish & Shellfish Immunology, 32, 273–283. doi: 10.1016/j.fsi.2011.11.016

Luo, Q., Tang, Q., Deng, L., Duan, Q., Zhang, R. (2023). A new cavefish of Sinocyclocheilus (Teleostei: Cypriniformes: Cyprinidae) from the Nanpanjiang River in Guizhou, China. Journal of Fish Biology, n/a. doi: 10.1111/jfb.15490

Mao, J., Huang, X., Sun, H., Jin, X., Guan, W., Xie, J., Wang, Y., Wang, X., Yin, D., Hao, Z., Tian, Y., Song, J., Ding, J., Chang, Y. (2022a). Transcriptome analysis provides insight into adaptive mechanisms of scallops under environmental stress. Frontiers in Marine Science, 9. doi: 10.3389/fmars.2022.971796

Mao, T., Liu, Y., Vasconcellos, M. M., Pie, M. R., Ellepola, G., Fu, C., Yang, J., Meegaskumbura, M. (2022b). Evolving in the darkness: phylogenomics of *Sinocyclocheilus* cavefishes highlights recent diversification and cryptic diversity. Molecular Phylogenetics and Evolution, 168, 107400. doi: 10.1016/j.ympev.2022.107400

Mao, T. R., Liu, Y. W., Meegaskumbura, M., Yang, J., Ellepola, G., Senevirathne, G., Fu, C., Gross, J. B., Pie, M. R. (2021). Evolution in *Sinocyclocheilus* cavefish is marked by rate shifts, reversals and origin of novel traits. Bmc Ecology and Evolution, 21, 45. doi: 10.1186/s12862-021-01776-y

Mayer, A., Mora, T., Rivoire, O., Walczak, A. M. (2016). Diversity of immune strategies explained by adaptation to pathogen statistics. Proc Natl Acad Sci U S A, 113, 8630–8635. doi: 10.1073/pnas.1600663113

Mccormick, M. I., Larson, J. K. (2008). Effect of hunger on the response to, and the production of, chemical alarm cues in a coral reef fish. Animal Behaviour, 75, 1973–1980. doi:10.1016/j.anbehav.2007.12.007

Mcgaugh, S. E., Gross, J. B., Aken, B., Blin, M., Borowsky, R., Chalopin, D., Hinaux, H., Jeffery, W. R., Keene, A., Ma, L., Minx, P., Murphy, D., O’quin, K. E., Retaux, S., Rohner, N., Searle, S. M., Stahl, B. A., Tabin, C., Volff, J. N., Yoshizawa, M., Warren, W. C. (2014). The cavefish genome reveals candidate genes for eye loss. Nat Commun, 5, 5307. doi: 10.1038/ncomms6307

Meng, F., Braasch, I., Phillips, J. B., Lin, X., Titus, T., Zhang, C., Postlethwait, J. H. (2013). Evolution of the eye transcriptome under constant darkness in *Sinocyclocheilus* cavefish. Molecular Biology and Evolution, 30, 1527–1543. doi: 10.1093/molbev/mst079

Meng, F. W., Zhao, Y. H., Titus, T., Zhang, C. G., Postlethwait, J. H. (2018). Brain of the blind: transcriptomics of the golden-line cavefish brain. Current Zoology, 64, 765–773. doi: 10.1093/cz/zoy005

Mohanty, B. R., Sahoo, P. K. (2010). Immune responses and expression profiles of some immune-related genes in Indian major carp, Labeo rohita to Edwardsiella tarda infection. Fish & Shellfish Immunology, 28, 613–621. doi: 10.1016/j.fsi.2009.12.025

Moran, D., Softley, R., Warrant, E. J. (2014). Eyeless Mexican cavefish save energy by eliminating the circadian rhythm in metabolism. PLoS One, 9, e107877. doi: 10.1371/journal.pone.0107877

Moran, D., Softley, R., Warrant, E. J. (2015). The energetic cost of vision and the evolution of eyeless Mexican cavefish. Science advances, 1, e1500363. doi: 10.1126/sciadv.1500363

Moran, R. L., Richards, E. J., Ornelas-García, C. P., Gross, J. B., Donny, A., Wiese, J., Keene, A. C., Kowalko, J. E., Rohner, N., Mcgaugh, S. E. (2023). Selection-driven trait loss in independently evolved cavefish populations. Nature Communications, 14, 2557. doi: 10.1038/s41467-023-37909-8

Morikawa, T., Takubo, K. (2016). Hypoxia regulates the hematopoietic stem cell niche. Pflügers Archiv-European Journal of Physiology, 468, 13–22. doi: 10.1007/s00424-015-1743-z

Nilsson Sköld, H., Aspengren, S., Wallin, M. (2013). Rapid color change in fish and amphibians– function, regulation, and emerging applications. Pigment cell & melanoma research, 26, 29–38. doi: 10.1111/pcmr.12040

Okar, D. A., Lange, A. J. (1999). Fructose-2, 6-bisphosphate and control of carbohydrate metabolism in eukaryotes. Biofactors, 10, 1–14. doi:10.1002/biof.5520100101

Peuß, R., Box, A. C., Chen, S., Wang, Y., Tsuchiya, D., Persons, J. L., Kenzior, A., Maldonado, E., Krishnan, J., Scharsack, J. P. (2020). Adaptation to low parasite abundance affects immune investment and immunopathological responses of cavefish. Nature ecology & evolution, 4, 1416–1430. doi: 10.1038/s41559-020-1234-2

Pham, L., Komalavilas, P., Eddie, A. M., Thayer, T. E., Greenwood, D. L., Liu, K. H., Weinberg, J., Patterson, A., Fessel, J. P., Boyd, K. L., Schafer, J. C., Kuck, J. L., Shaver, A. C., Flaherty, D. K., Matlock, B. K., Wijers, C. D. M., Serezani, C. H., Jones, D. P., Brittain, E. L., Rathmell, J. C., Noto, M. J. (2022). Neutrophil trafficking to the site of infection requires Cpt1a-dependent fatty acid β-oxidation. Commun Biol, 5, 1366. doi: 10.1038/s42003-022-04339-z

Policarpo, M., Fumey, J., Lafargeas, P., Naquin, D., Thermes, C., Naville, M., Dechaud, C., Volff, J.N., Cabau, C., Klopp, C. (2021). Contrasting gene decay in subterranean vertebrates: insights from cavefishes and fossorial mammals. Molecular Biology and Evolution, 38, 589–605. doi: 10.1093/molbev/msaa249

Policarpo, M., Fumey, J., Lafargeas, P., Naquin, D., Thermes, C., Naville, M., Dechaud, C., Volff, J.N., Cabau, C., Klopp, C., Møller, P. R., Bernatchez, L., García-Machado, E., Rétaux, S., Casane, D. (2020). Contrasting Gene Decay in Subterranean Vertebrates: Insights from Cavefishes and Fossorial Mammals. Molecular Biology and Evolution, 38, 589–605. doi: 10.1093/molbev/msaa249

Protas, M., Jeffery, W. R. (2012). Evolution and development in cave animals: from fish to crustaceans. Wiley Interdiscip Rev Dev Biol, 1, 823–845. doi: 10.1002/wdev.61

Rambaut, A. (2009). FigTree. Tree figure drawing tool: http://tree.bio.ed.ac.uk/software/figtree/

Qi, D., Chao, Y., Wu, R., Xia, M., Chen, Q., Zheng, Z. (2018). Transcriptome Analysis Provides Insights Into the Adaptive Responses to Hypoxia of a Schizothoracine Fish (Gymnocypris eckloni). Front Physiol, 9, 1326. doi: 10.3389/fphys.2018.01326

Randhawa, M., Sangar, V., Tucker-Samaras, S., Southall, M. (2014). Metabolic signature of sun exposed skin suggests catabolic pathway overweighs anabolic pathway. PLoS One, 9, e90367. doi:10.1371/journal.pone.0090367

Reynolds, T. B. (2009). Strategies for acquiring the phospholipid metabolite inositol in pathogenic bacteria, fungi and protozoa: making it and taking it. Microbiology, 155, 1386. doi:10.1099/mic.0.025718-0

Riddle, M. R., Aspiras, A. C., Gaudenz, K., Peuß, R., Sung, J. Y., Martineau, B., Peavey, M., Box, A. C., Tabin, J. A., Mcgaugh, S. (2018). Insulin resistance in cavefish as an adaptation to a nutrient-limited environment. Nature, 555, 647–651. doi: 10.1038/nature26136

Romero, A., Green, S. M. (2005). The end of regressive evolution: examining and interpreting the evidence from cave fishes. Journal of Fish Biology, 67, 3–32. doi: 10.1111/j.0022-1112.2005.00776.x

Romero, A., Paulson, K. M. (2001). It’sa wonderful hypogean life: a guide to the troglomorphic fishes of the world. The biology of hypogean fishes, 13–41. doi: 10.1023/A:1011844404235

Rzepka, Z., Buszman, E., Beberok, A., Wrześniok, D. (2016). From tyrosine to melanin: Signaling pathways and factors regulating melanogenesis. Postepy Hig Med Dosw (Online), 70, 695–708. doi: 10.5604/17322693.1208033

Sato, Y., Tsurumi, T. (2013). Genome guardian p53 and viral infections. Reviews in Medical Virology, 23, 213–220. doi: 10.1002/rmv.1738

Schäfer, B. W., Heizmann, C. W. (1996). The S100 family of EF-hand calcium-binding proteins: functions and pathology. Trends Biochem Sci, 21, 134–140. doi: 10.1016/s0968-0004(96)80167-8

Scharsack, J. P., Kalbe, M., Harrod, C., Rauch, G. (2007). Habitat-specific adaptation of immune responses of stickleback (Gasterosteus aculeatus) lake and river ecotypes. Proceedings of the Royal Society B: Biological Sciences, 274, 1523–1532. doi: 10.1098/rspb.2007.0210

Simon, V., Elleboode, R., Mahé, K., Legendre, L., Ornelas-Garcia, P., Espinasa, L., Rétaux, S. (2017). Comparing growth in surface and cave morphs of the species *Astyanax mexicanus*: insights from scales. EvoDevo, 8, 1–13. doi: 10.1186/s13227-017-0086-6

Soares, D., Niemiller, M. L. (2020). Extreme adaptation in caves. The Anatomical Record, 303, 15–23. doi:10.1002/ar.24044

Solaini, G., Baracca, A., Lenaz, G., Sgarbi, G. (2010). Hypoxia and mitochondrial oxidative metabolism. Biochimica et Biophysica Acta (BBA) – Bioenergetics, 1797, 1171–1177. doi: 10.1016/j.bbabio.2010.02.011

Stahl, B. A., Gross, J. B. (2015). Alterations in Mc1r gene expression are associated with regressive pigmentation in *Astyanax* cavefish. Dev Genes Evol, 225, 367–375. doi: 10.1007/s00427-015-0517-0

Stahl, B. A., Gross, J. B. (2017). A comparative transcriptomic analysis of development in two *Astyanax* cavefish populations. Journal of Experimental Zoology Part B: Molecular and Developmental Evolution, 328, 515–532. doi: 10.1002/jez.b.22749

Staudigl, M., Gersting, S. W., Danecka, M. K., Messing, D. D., Woidy, M., Pinkas, D., Kemter, K. F., Blau, N., Muntau, A. C. (2011). The interplay between genotype, metabolic state and cofactor treatment governs phenylalanine hydroxylase function and drug response. Human Molecular Genetics, 20, 2628–2641. doi: 10.1093/hmg/ddr165

Sun, Y., Liu, W.Z., Liu, T., Feng, X., Yang, N., Zhou, H.F. (2015). Signaling pathway of MAPK/ERK in cell proliferation, differentiation, migration, senescence and apoptosis. Journal of Receptors and Signal Transduction, 35, 600–604. doi: 10.3109/10799893.2015.1030412

Thiergart, T., Durán, P., Ellis, T., Vannier, N., Garrido-Oter, R., Kemen, E., Roux, F., Alonso-Blanco, C., Ågren, J., Schulze-Lefert, P., Hacquard, S. (2020). Root microbiota assembly and adaptive differentiation among European Arabidopsis populations. Nature ecology & evolution, 4, 122–131. doi: 10.1038/s41559-019-1063-3

Tong, C., Li, M., Tang, Y., Zhao, K. (2021). Genomic Signature of Shifts in Selection and Alkaline Adaptation in Highland Fish. Genome Biology and Evolution, 13. doi: 10.1093/gbe/evab086

Torres-Paz, J., Hyacinthe, C., Pierre, C., Rétaux, S. (2018). Towards an integrated approach to understand Mexican cavefish evolution. Biol Lett, 14. doi: 10.1098/rsbl.2018.0101

Voskarides, K., Koutsofti, C., Pozova, M. (2022). TP53 Mutant Versus Wild-Type Zebrafish Larvae Under Starvation Stress: Larvae Can Live Up to 17 Days Post-Fertilization Without Food. Zebrafish, 19, 49–55. doi: 10.1089/zeb.2022.0003

Wen, H., Luo, T., Wang, Y., Wang, S., Liu, T., Xiao, N., Zhou, J. (2022). Molecular phylogeny and historical biogeography of the cave fish genus *Sinocyclocheilus* (Cypriniformes: Cyprinidae) in southwest China. Integrative Zoology, 17, 311–325. doi: 10.1111/1749-4877.12624

Xiao, W., Zou, Z., Li, D., Zhu, J., Yue, Y., Yang, H. (2020). Effect of dietary phenylalanine level on growth performance, body composition, and biochemical parameters in plasma of juvenile hybrid tilapia, Oreochromis niloticus × Oreochromis aureus. Journal of the World Aquaculture Society, 51, 437–451. doi: 10.1111/jwas.12641

Xie, C., Mao, X. Z., Huang, J. J., Ding, Y., Wu, J. M., Dong, S., Kong, L., Gao, G., Li, C.Y., Wei, L. P. (2011). KOBAS 2.0: a web server for annotation and identification of enriched pathways and diseases. Nucleic Acids Research, 39, W316–W322. doi: 10.1093/nar/gkr483

Xiong, S. (2021). How cavefish gain high body fat to adapt to food scarcity: Open University (United Kingdom).

Xu, C., Luo, T., Zhou, J.-J., Wu, L., Zhao, X.-R., Yang, H.-F., Xiao, N., Zhou, J. (2023). Sinocyclocheilus longicornus (Cypriniformes, Cyprinidae), a new species of microphthalmic hypogean fish from Guizhou, Southwest China. ZooKeys, 1141, 1–28. doi: 10.3897/zookeys.1141.91501

Yang, J., Chen, X., Bai, J., Fang, D., Qiu, Y., Jiang, W., Yuan, H., Bian, C., Lu, J., He, S., Pan, X., Zhang, Y., Wang, X., You, X., Wang, Y., Sun, Y., Mao, D., Liu, Y., Fan, G., Zhang, H., Chen, X., Zhang, X., Zheng, L., Wang, J., Cheng, L., Chen, J., Ruan, Z., Li, J., Yu, H., Peng, C., Ma, X., Xu, J., He, Y., Xu, Z., Xu, P., Wang, J., Yang, H., Wang, J., Whitten, T., Xu, X., Shi, Q. (2016). The *Sinocyclocheilus* cavefish genome provides insights into cave adaptation. Bmc Biology, 14, 1–13. doi: 10.1186/s12915-015-0223-4

Yang, Z. (2007). PAML 4: phylogenetic analysis by maximum likelihood. Molecular Biology and Evolution, 24, 1586–1591. doi: 10.1093/molbev/msm088

Yeung, K. Y., Ruzzo, W. L. (2001). Principal component analysis for clustering gene expression data. Bioinformatics, 17, 763–774. doi: 10.1093/bioinformatics/17.9.763

Yoshizawa, M., Goricki, S., Soares, D., Jeffery, W. R. (2010). Evolution of a behavioral shift mediated by superficial neuromasts helps cavefish find food in darkness. Curr Biol, 20, 1631–1636. doi: 10.1016/j.cub.2010.07.017

Yoshizawa, M., Jeffery, W. R., Van Netten, S. M., Mchenry, M. J. (2014). The sensitivity of lateral line receptors and their role in the behavior of Mexican blind cavefish (*Astyanax mexicanus*). Journal of Experimental Biology, 217, 886–895. doi:10.1242/jeb.094599

Young, M. D., Wakefield, M. J., Smyth, G. K., Oshlack, A. (2010). Gene ontology analysis for RNA-seq: accounting for selection bias. Genome Biol, 11, R14. doi: 10.1186/gb-2010-11-2-r14

Yu, S., Rao, S., Reddy, J. K. (2003). Peroxisome proliferator-activated receptors, fatty acid oxidation, steatohepatitis and hepatocarcinogenesis. Curr Mol Med, 3, 561–572. doi: 10.2174/1566524033479537

Zhang, Y., Liang, S., He, J., Bai, Y., Niu, Y., Tang, X., Li, D., Chen, Q. (2015). Oxidative stress and antioxidant status in a lizard Phrynocephalus vlangalii at different altitudes or acclimated to hypoxia. Comparative Biochemistry and Physiology Part A: Molecular & Integrative Physiology, 190, 9–14. doi: 10.1016/j.cbpa.2015.08.013

Zhao, Q., Shao, F., Li, Y., Yi, S. V., Peng, Z. (2022). Novel genome sequence of Chinese cavefish (Triplophysa rosa) reveals pervasive relaxation of natural selection in cavefish genomes. Molecular ecology, 31, 5831–5845. doi: 10.1111/mec.16700

Zhao, Y., Huang, X., Ding, T. W., Gong, Z. (2016). Enhanced angiogenesis, hypoxia and neutrophil recruitment during Myc-induced liver tumorigenesis in *zebrafish*. Scientific reports, 6, 31952. doi: 10.1038/srep31952

Zhao, Y., Huang, Z., Huang, J., Zhang, C., Meng, F. (2021). Phylogenetic analysis and expression differences of eye-related genes in cavefish genus *Sinocyclocheilus*. Integr Zool, 16, 354–367. doi: 10.1111/1749-4877.12466

Zhao, Y. C., Chen, H. Y., Li, C. Q., Chen, S. Y., Xiao, H. (2020). Comparative Transcriptomics Reveals the Molecular Genetic Basis of Cave Adaptability in *Sinocyclocheilus* Fish Species. Frontiers in Ecology and Evolution, 8. doi: 10.3389/fevo.2020.589039

Zhao, Y. H., Zhang, C. G. (2009). Endemic fishes of Sinocyclocheilus (Cypriniformes: Cyprinidae) in China-species diversity, cave adaptation, systematics and zoogeograph, Beijing: Science Press.

Zhou, J., Luo, J., Zeng, Y., Xu, L. (2021). Characterization of PAH Gene Mutations and Analysis of Genotype-Phenotype Correlation in Patients with Phenylalanine Hydroxylase Deficiency from Fujian Province, Southeastern China. Molecular biology reports, 49, 10409–10419. doi: 10.21203/rs.3.rs-1096859/v1

Zhou, Q. L., Wang, L. Y., Zhao, X. L., Yang, Y. S., Ma, Q., Chen, G. (2022a). Effects of salinity acclimation on histological characteristics and miRNA expression profiles of scales in juvenile rainbow trout (Oncorhynchus mykiss). Bmc Genomics, 23, 300. doi: 10.1186/s12864-022-08531-7

Zhou, S., Rajput, A. P., Mao, T., Liu, Y., Ellepola, G., Herath, J., Yang, J., Meegaskumbura, M. (2022b). Adapting to Novel Environments Together: Evolutionary and Ecological Correlates of the Bacterial Microbiome of the World’s Largest Cavefish Diversification (Cyprinidae, Sinocyclocheilus). Frontiers in Microbiology, 13, 823254. doi: 10.3389/fmicb.2022.823254

Zhu, D., Zhang, C., Liu, P., Jawad, L. A. (2019). Comparison of the morphology, structures and mechanical properties of teleost fish scales collected from New Zealand. Journal of Bionic Engineering, 16, 328–336. doi: 10.1007/s42235-019-0028-1

