## Supplementary Figure S1 for "Habitat correlates of cave-dwelling: A radiation-scale analysis of skin traits and comparative transcriptomics of the *Sinocyclocheilus* cavefish"

| Pictures/Species | Digital camera | Leica M165FC stereoscopic microscope | Body length  (n ± σ mm) | | Colour morphology, Body colour RGB value: (n, n, n) and Black blotches: (n %) | | Scale morphology | | Average area of lateral line scales  (n ± σ mm^2^) | | Number of lateral line scales  (n ± σ) | | Reference | |
| --- | --- | --- | --- | --- | --- | --- | --- | --- | --- | --- | --- | --- | --- | --- |
| S. *furcodorsalis*   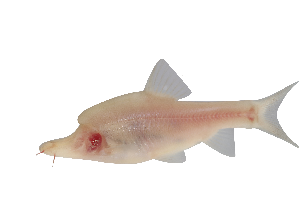 | 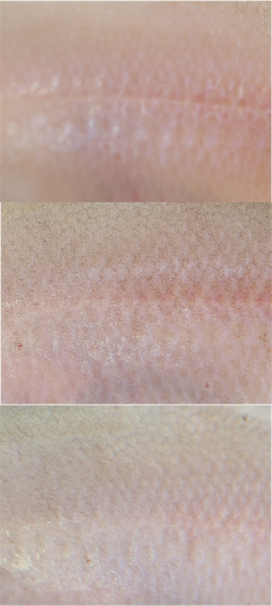 | 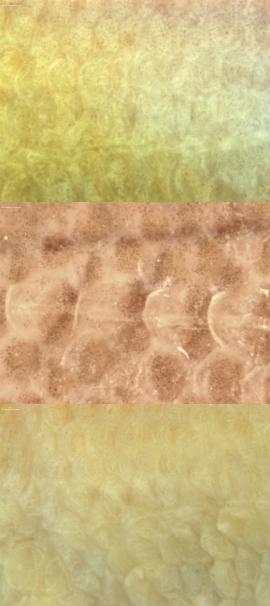 | 97.419±3.266 | | Pink,translucent (188,163,155);  The specimen is creamy yellow with small number of tiny light gray spots on the body (0.00). | | Incompletely covered with big membranous scales, some of the scales on the sides of the body buried under the skin and disappeared;  Lateral line scales slightly larger than upper and lower body scales. | | 2.632 ±0.285 | | 38±4 | | this study | |
| *S. tianeensis*  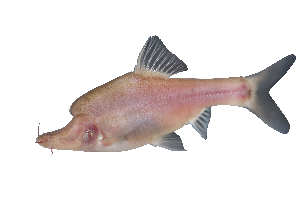 | 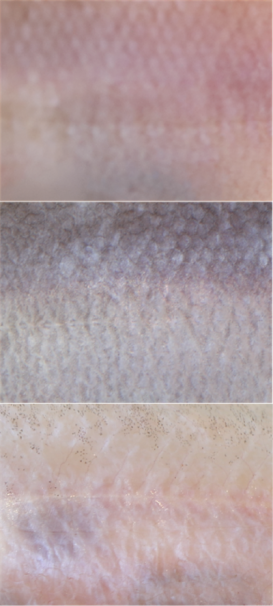 | 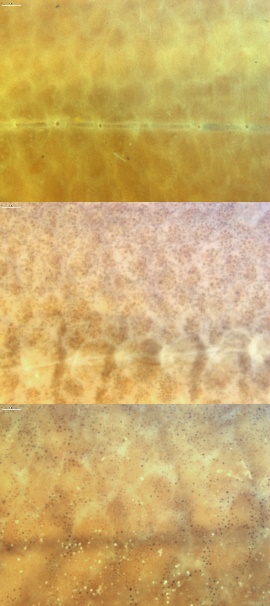 | 98.541±12.115 | | Pink, translucent (174,155,157);  The specimen is creamy yellow with small number of tiny light gray spots on the body (0.00). | | Incompletely covered with big membranous scales, some of the scales on the sides of the body buried under the skin and disappeared;  Lateral line scales larger than upper and lower body scales. | | 2.022 ±1.319 | | 37±2 | | this study | |
| *S. xunlensis*  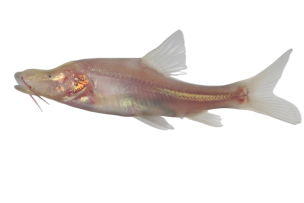 | 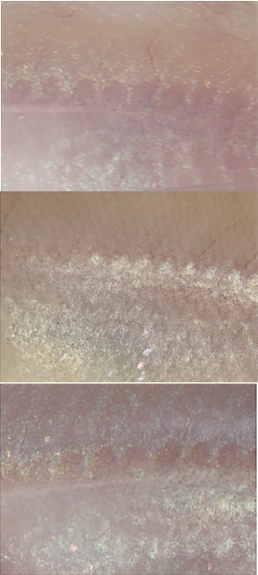 | 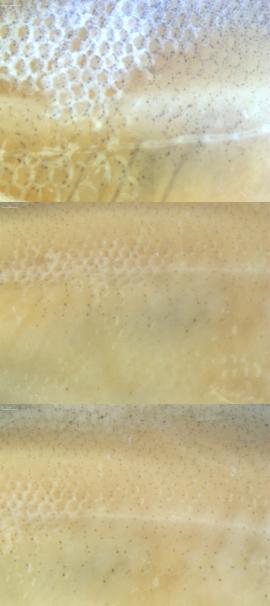 | 47.854±6.676 | | Pink, translucent (173,148,140);  The specimen is creamy yellow (0.00). | | Completely covered with small membranous scales; Lateral line scales similar in size to body scales. | | 0.654 ±0.367 | | 43±4 | | this study | |
| *S. cyphotergous*  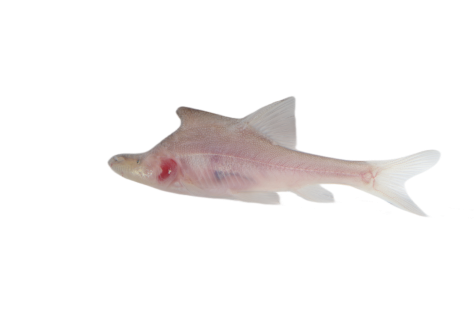 | 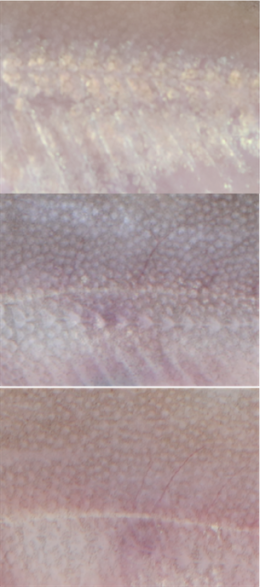 | 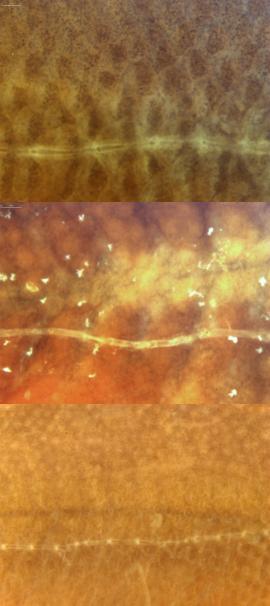 | 85.301±7.283 | | Gray, translucent (174,155,157);  The specimen is yellowish brown with a slightly darker dorsal colour, with large number of tiny black gray spots on the body (2.12). | | Incompletely covered with small membranous scales, body scales buried under the skin and some ventral scales disappeared ;  Lateral line scales slightly larger than upper and lower body scales. | | 1.883 ±0.676 | | 52±10 | | this study | |
| *S. mashanensis*  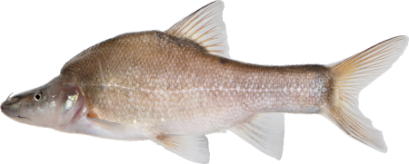 | 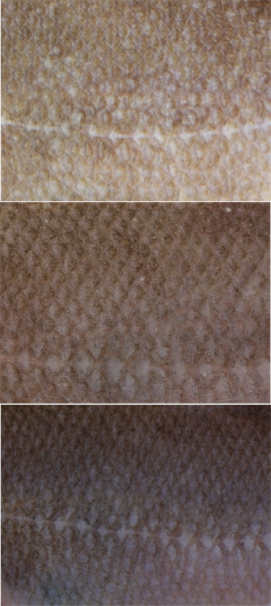 | 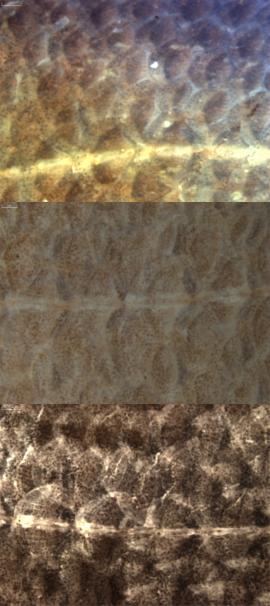 | 95.055±11.000 | | Gray, translucent (151, 127, 127);  The specimen is yellowish brown with a slightly darker dorsal colour, with large number of tiny black spots on the body (18.05). | | Completely covered with big scales;  Lateral line scales slightly larger than upper and lower body scales. | | 3.742 ±1.357 | | 49±3 | | this study | |
| *S. zhenfengensis*  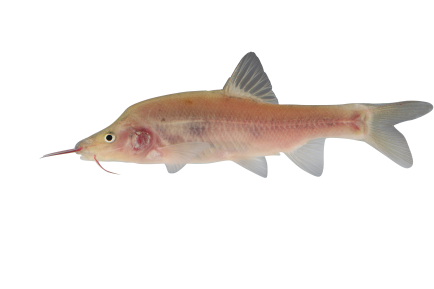 | 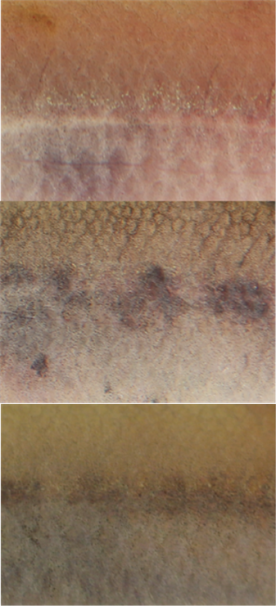 | 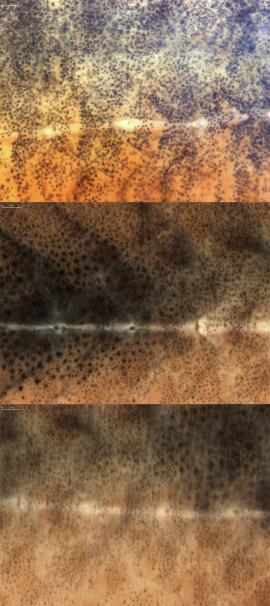 | 83.621±6.014 | | Gray, translucent with grey spots (145,116,93);  The specimen is yellowish brown with a darker dorsal colour, with big black spots on the body (7.93). | | Completely covered with big scales;  Lateral line scales slightly larger than upper and lower body scales. | | 2.934 ±0.954 | | 43±4 | | this study | |
| *S. guanyangensis*  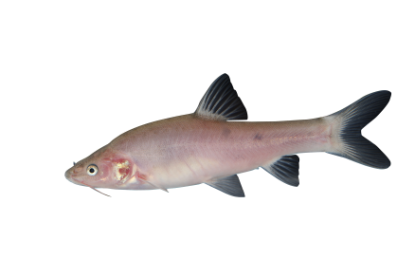 | 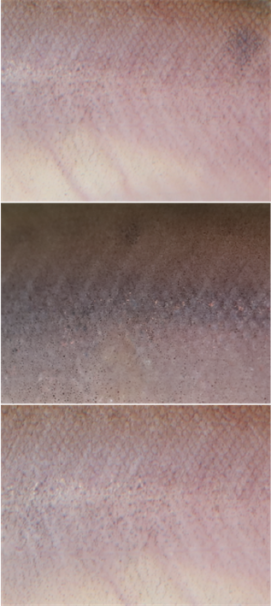 | 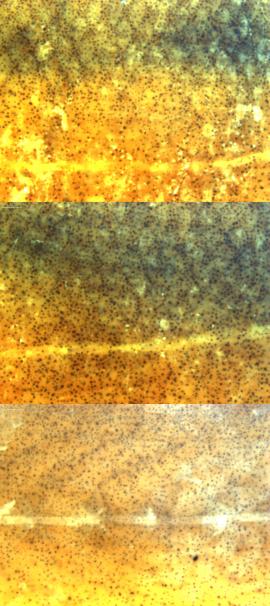 | 106.265±25.033 | | Light grayish white with black spots (133, 112, 98);  The specimen is yellowish brown with a slightly darker dorsal colour, with big black spots on the back (10.15). | | Completely covered with small scales;  Lateral line scales similar in size to body scales. | | 1.157 ±0.266 | | 57±4 | | this study | |
| *S. guilinensis*  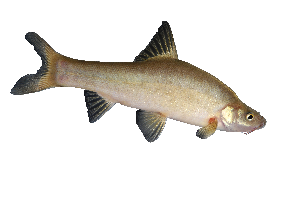 | 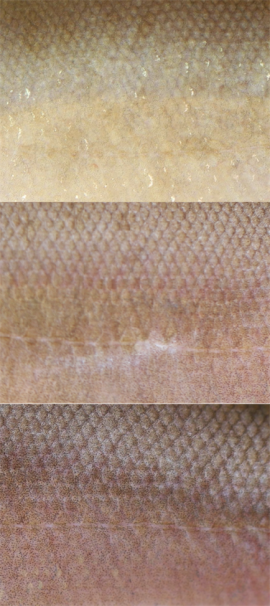 | 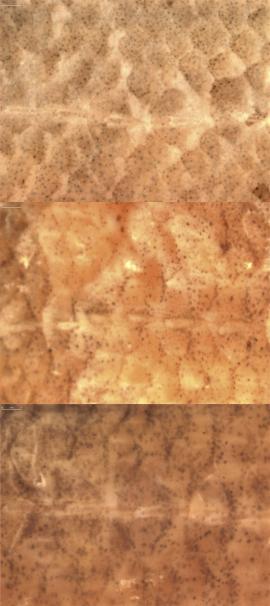 | 106.831±14.773 | | Golden yellow without black spots (165,140,121);  The specimen is creamy yellow with a slightly darker dorsal colour, with small number of tiny light gray spots on the body (12.35). | | Completely covered with big scales;  Lateral line scales larger than upper and lower body scales. | | 2.000 ±0.615 | | 50±8 | | this study | |
| *S.angustiporus*  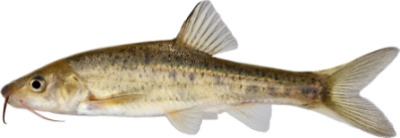 | 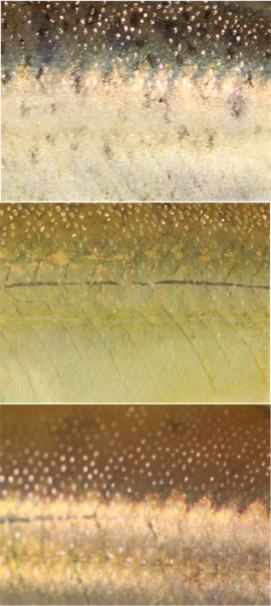 | 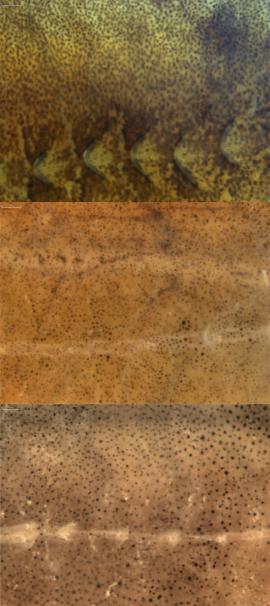 | 97.767±32.681 | | Yellow, small number of black spots (173,150,105);  The specimen is yellowish brown; with big black spots on the body (9.93). | | Incompletely covered with tiny scales, body scales mostly buried under the skin and disappeared;  Lateral line scales only, with most of the scales above and below the lateral line disappeared. | | 1.707 ±0.675 | | 72±11 | | this study | |
| *S. qiubeiensis*  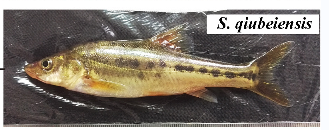 | 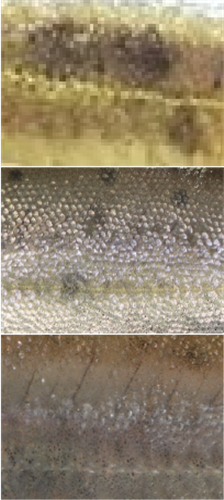 | 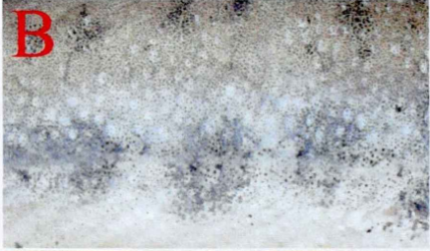 | 99.692±3.058 | | Yellow, small number of black spots (142,127,104);  The specimen is yellowish brown; with with big black spots on the body (26.39). | | Completely covered with tiny scales, some body scales hidden under the skin and disappeared;  Lateral line scales larger than upper and lower body scales. | | 0.605 ±0.277 | | 78±3 | | Li et al., 2020 | |
| *S. oxycephalus*  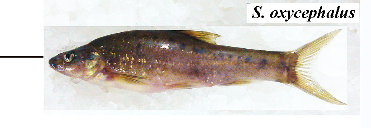 | 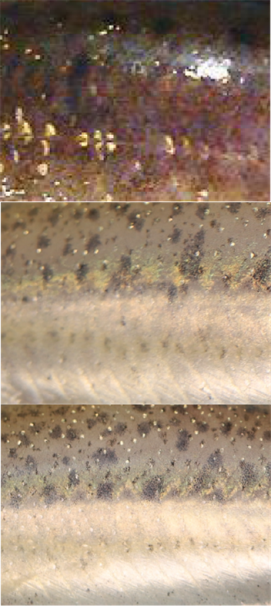 | 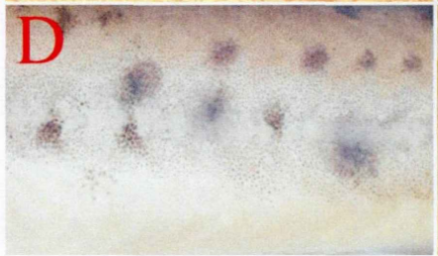 | 110.122±2.979 | | Charcoal grey, small number of black spots (142,116,90);  The specimen is yellowish brown with with big black spots on the body (30.32). | | Incompletely covered with tiny scales, body scales mostly buried under the skin and disappeared;  Lateral line scales larger than upper and lower body scales. | | 0.165 ±0.106 | | 70±3 | | Li et al., 2020 | |
| *S. tianlinensis*  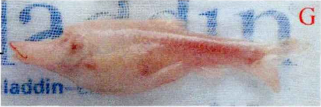 | 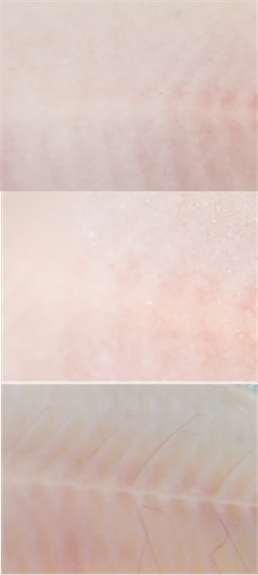 | 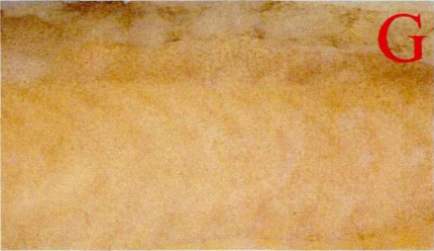 | 70.954±1.313 | | Pink, translucent (195,181,177);  The specimen is creamy yellow with small number of tiny light gray spots on the body (2.08). | | Body scales and lateral line scales absent. | | 0 | | 0 | | Li et al., 2020 | |
| *S. anshuiensis*  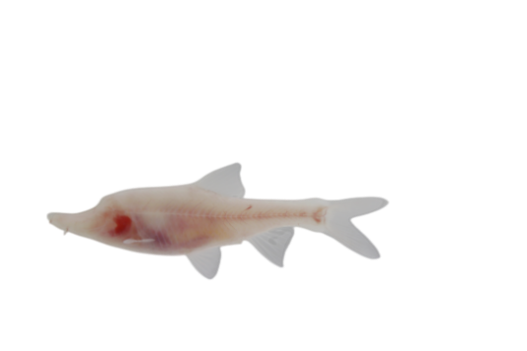 | 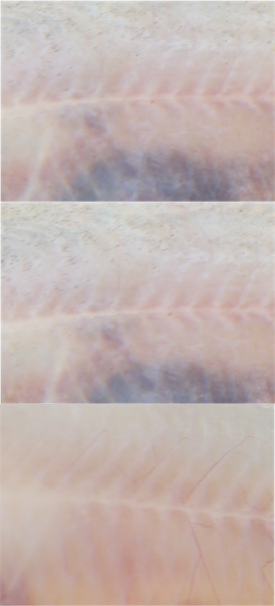 | 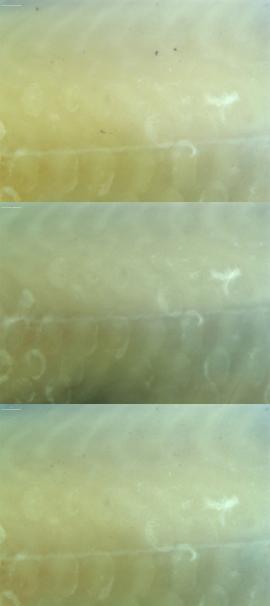 | 70.216±2.356 | | Pink, translucence (191,179,177);  The specimen is creamy yellow (0.00). | | Incompletely covered with samll membranous scales;  Lateral line scales larger than upper and lower body scales. | | 1.816 ±0.273 | | 35±1 | | this study | |
| *S. grahami*  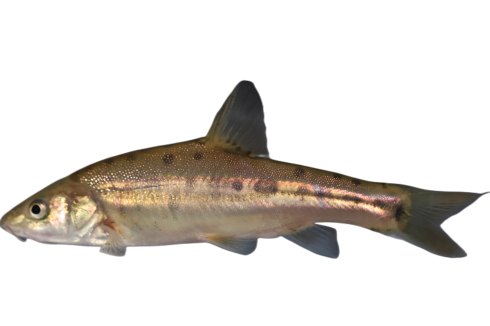 | 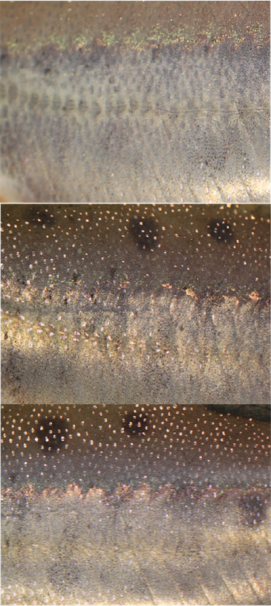 | 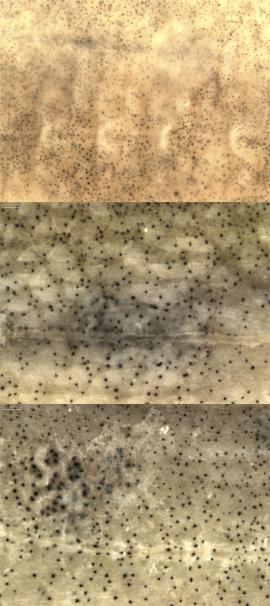 | | 117.712±1.273 | | Golden yellow, few black spots (138,122,108).  The specimen is yellowish brown; with big black spots on the body (34.09). | | Incompletely covered with tiny scales, body scales mostly buried under the skin;  Lateral line scales larger than upper and lower body scales. | | 0.720 ±0.396 | | 72±2 | | this study |

**Supplementary Figure S2. The skin morphology of each *Sinocyclocheilus* speciesindicatingtheir photos, body colour and pigmentation, characteristic of the number of scales. (“n” represents the mean and “σ” represents the standard deviation).**
