## Supplementary Table S1 for "Habitat correlates of cave-dwelling: A radiation-scale analysis of skin traits and comparative transcriptomics of the *Sinocyclocheilus* cavefish"

**Supplementary Table S1. Sampling details of 14 *Sinocyclocheilus* species indicating their clades, location, habitats,** **eye-types and source of individuals.**

| **Species** | **clades** | **Latitude**  **(N)** | **Longitude (E)** | **Habitats** | **Eye morphology** | **Reference** |
| --- | --- | --- | --- | --- | --- | --- |
| S. *furcodorsalis* | B | 24.9506 | 107.0533 | Stygobitic | Eyeless | this study |
| *S. tianlinsis* | B | 24.4900 | 106.3600 | Stygobitic | Eyeless | Li et al., 2020 |
| *S. tianeensis* | B | 24.8800 | 107.2000 | Stygobitic | Eyeless | this study |
| *S. anshuiensis* | B | 24.3758 | 106.7556 | Stygobitic | Eyeless | Yang et al., 2016 |
| *S. guanyangensis* | A | 25.2283 | 110.9292 | Stygophilic | Normal-eyed | this study |
| *S. mashanensis* | B | 23.8544 | 108.5078 | Stygophilic | Micro-eyed | this study |
| *S. xunlensis* | C | 25.4000 | 108.2000 | Stygophilic | Micro-eyed | this study |
| *S. cyphotergous* | C | 25.5904 | 106.6813 | Stygophilic | Micro-eyed | this study |
| *S. guilinensis* | A | 25.1075 | 110.2649 | Surface | Normal-eyed | this study |
| *S. zhenfengensis* | B | 25.4679 | 105.6456 | Stygophilic | Normal-eyed | this study |
| *S. oxycephalus* | D | 24.7696 | 103.3122 | Surface | Normal-eyed | Li et al., 2020 |
| *S. angustiporus* | D | 25.1408 | 104.1167 | Surface | Normal-eyed | this study |
| *S. qiubeiensis* | D | 24.2779 | 104.0403 | Surface | Normal-eyed | Li et al., 2020 |
| *S. grahami* | D | 24.9646 | 102.6663 | Surface | Normal-eyed | Yang et al., 2016 |
